# A co-ordinated transcriptional programme in the maternal liver supplies LC-PUFAs to the conceptus using phospholipids

**DOI:** 10.1101/2023.06.23.546226

**Authors:** Risha Amarsi, Samuel Furse, Mary AM Cleaton, Sarah Maurel, Alice L Mitchell, Anne C. Ferguson-Smith, Nicolas Cenac, Catherine Williamson, Albert Koulman, Marika Charalambous

**Affiliations:** Department of Medical and Molecular Genetics, Faculty of Life Sciences and Medicine, King's College London, London SE19RT, UK; Biological chemistry group, Jodrell laboratory, Royal Botanic Gardens Kew, Kew Road, Richmond, Surrey, TW9 3DS, United Kingdom; Core Metabolomics and Lipidomics Laboratory, Wellcome-MRC Institute of Metabolic Science, University of Cambridge, Addenbrooke’s Treatment Centre, Keith Day Road, Cambridge, CB2 0QQ, United Kingdom; Department of Genetics, Downing Street, University of Cambridge, Cambridge CB2 3EH, UK; IRSD, Université de Toulouse-Paul Sabatier, INSERM, INRAe, ENVT, UPS, Toulouse, France; Department of Women and Children's Health, King's College London, Guy's Campus, London, UK

## Abstract

Essential fatty acids (EFAs) and their derivatives, the long and very long chain polyunsaturated fatty acids (LC-PUFAs), are preferentially transported by the mother to the fetus. Failure to supply EFAs is strongly linked with stillbirth, fetal growth restriction, and impaired neurodevelopmental outcomes. However, dietary supplementation during pregnancy is unable to simply reverse these outcomes, suggesting imperfectly understood interactions between dietary EFA intake and the molecular mechanisms of maternal supply. Here we combine untargeted lipidomics with transcriptional profiling of healthy and genetically-manipulated murine models to understand the maternal adaptations required to provide LC-PUFAs to the developing fetus. We discovered a late pregnancy-specific, selective activation of the Liver X Receptor signalling pathway which dramatically increases maternal supply of LC-PUFAs within circulating phospholipids. Crucially, genetic ablation of this pathway in the mother reduced LC-PUFA accumulation by the fetus. Overall our work suggests new molecular strategies for improving maternal-fetal transfer of these important lipids.

## INTRODUCTION

During a normal pregnancy, maternal lipid metabolism undergoes striking adaptations to meet the nutritional demands of the developing fetus. Anabolic pathways in early pregnancy promote the net accumulation of nutrients within maternal tissues, primarily in the form of triglycerides (1). A mid-gestation catabolic switch then drives the breakdown of nutrient stores, ensuring the circulating availability of fatty acids and glucose for fetal uptake. These dynamically regulated shifts in maternal nutrient allocation are vital for a healthy pregnancy, as poor adaptations, commonly seen in obese pregnancy, can profoundly disrupt the development and long-term metabolic health of the child (2).

Rodent studies have helped to characterise the anabolic and catabolic shifts in whole-body lipid metabolism (3). In a previous study we demonstrated that maternal plasma Delta-like homologue 1 (DLK1) levels are elevated in the catabolic phase of pregnancy, and that the major source of this protein was the conceptus (4). *Dlk1* is an imprinted gene expressed predominantly from the paternally-inherited allele (5,6). This mode of inheritance allowed us to independently examine the influence of *Dlk1* loss of function intrinsically in maternal tissues or as an endocrine factor produced by the fetus. We observed that loss of DLK1 in the fetus caused impairments in maternal fasting metabolism and lipoprotein production, whereas loss of DLK1 in the dam prevented the normal acquisition and release of her adipose tissue stores (4). This work led us to hypothesise that DLK1 is a key modulator of maternal fatty acid metabolism in pregnancy.

Fatty acids (FAs) are crucial components at every stage of fetal development, not just as an energy source, but also to provide building blocks and key signals for organogenesis (7). Of particular importance are the essential fatty acids (EFAs), linoleic acid (LA, 18:2n-6) and alpha-linolenic acid (ALA, 18:3n-3), which can only be obtained from dietary sources. Through elongation/desaturation reactions, LA is used to synthesise omega-6 (n-6) long chain polyunsaturated fatty acids (LC-PUFA) like arachidonic acid (ARA, 20:4n-6), and ALA is converted into omega-3 (n-3) LC-PUFAs which include eicosapentaenoic acid (EPA, 20:5n-3) and docosahexaenoic acid (DHA, 22:6n-3), however only a small amount of nutrient requirement is thought to be met by this route in the non-pregnant state (8). Both n-6 and n-3 FAs are essential for fetal development, with roles in placentation, synthesis of membrane components and membrane fluidity, and the production of signalling molecules (9). They are especially important for brain growth and central nervous system development, since this organ contains 50-60% dry mass as lipid, ∼35% of which are LC-PUFAs (10). Fetal synthesis can only account for a small proportion of this demand, and instead maternal production/mobilisation and placental transfer is required to meet the ARA and DHA requirements for healthy development (11,12). However, the mechanism driving this selective “biomagnification” of LC-PUFAs between maternal and fetal compartments is not fully understood.

Although circulating adipose-derived non-esterified fatty acids (NEFAs) can be directly transported across the placenta, the majority are transferred to the maternal liver, a highly adaptive, yet poorly characterised metabolic tissue in pregnancy. Hepatic re-esterification of fatty acids into triglycerides and export into the circulation within very low-density lipoproteins (VLDL) underlies the characteristic rise in plasma triglycerides in late gestation and is considered the primary source of fatty acids for the fetus (7). In normal physiology, the liver modulates complex synthetic and catabolic pathways to regulate whole-body lipid dynamics. Since genetic polymorphisms within known hepatic lipid pathway genes drive at least 10% of population variation in plasma triglycerides (13,14), we propose that hepatic pathways are likely regulators of fatty acid allocation in pregnancy.

However, because of the focus on hepatic triglycerides as the main fetal source of LC-PUFAs, unbiased mechanistic investigations into lipid synthetic pathways in the maternal liver are scarce. In recent years, lipidomic investigations of pregnancy have provided a molecular insight into the fatty acid composition of lipids, with numerous phospholipid biomarkers reported for common metabolic complications of pregnancy (15–17). Phospholipids are well-reported to rise in the maternal circulation (18) and could represent a substantial physiological source of LC-PUFAs in plasma and tissues (19–21). These studies suggested to us that biomagnification pathways may extend beyond production of maternal circulating triglycerides and involve LC-PUFA incorporation into other types of lipid.

Here we profiled the liver and plasma lipidomes of virgin and pregnant mice as an untargeted discovery approach for lipid pathways in the maternal liver that are associated with maternal fatty acid provision to the conceptus. With the aid of our *Dlk1* model of disrupted lipid metabolism and targeted PUFA metabolite and transcriptional profiles, we aimed to delineate the major hepatic lipid pathways that underlie the fatty acid adaptations of late pregnancy. We show that i) ARA and DHA are enriched in the maternal liver and in the circulation selectively in the form of phospholipids; ii) Hepatic transcriptional pathways that synthesise LC-PUFAs from dietary intermediates and incorporate them into phospholipids are activated in late pregnancy; iii) This transcriptional programme is co-ordinately regulated by the liver X receptor (LXR) and modulated by maternal production of DLK1. Taken together we propose that biomagnification is achieved by a co-ordinated regulatory programme in the mother to supply LC-PUFAs to the conceptus in the phospholipid compartment.

## RESULTS

In the current study we used samples from a previously published cohort to investigate both maternal lipid metabolism in normal pregnancy, and perturbations to this process as a result of loss of DLK1 (4). Liver and plasma were collected from matched virgin and pregnant dams at 15.5 dpc in 8 groups with modified DLK1 in either the maternal tissue or in the plasma (derived from the conceptus), Figure 1A. We hypothesised that i) the abundance of specific lipid species would be co-ordinately modified in late pregnancy in the plasma and liver. To test this we compared 3 replicate groups of matched maternal genotype (Figure 1B). ii) Circulating DLK1 generated by the conceptus can influence maternal plasma-liver lipid abundance. Here we compared groups with and without a functional fetal *Dlk1* copy in matched genotype dams (Figure 1C, ‘Fetal Effect’. iii) Finally, we hypothesised that maternal *Dlk1* genotype would influence the abundance of maternal lipids, and tested this by comparing dams with matched fetal DLK1 production (Figure 1C, ‘Maternal Effect’).

**Figure 1.**
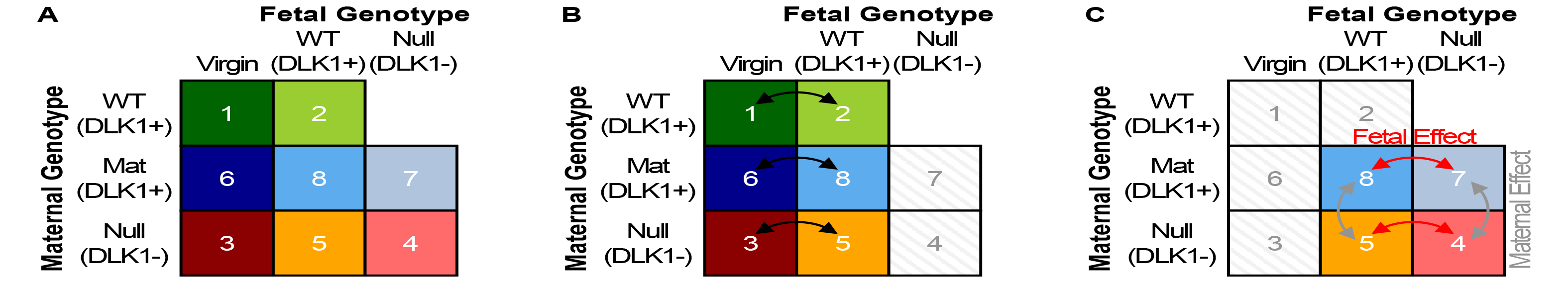
Study design. **(A)** We used liver and plasma samples from a previously published cohort with manipulations to the imprinted gene, *Dlk1*. The study included five pregnant groups at 15.5 dpc (groups 2, 4, 5, 7 and 8) and three age and genotype-matched virgin groups (groups 1, 3, and 6). The study included five pregnant groups at 15.5 dpc (groups 2, 4, 5, 7 and 8) and three age-matched virgin groups (groups 1, 3, and 6). Groups 1 and 2 were genetically unmodified females with normal DLK1 expression. Groups 6, 8 and 7 inherited a silent *Dlk1* deletion from their mother and so maintained a functional copy of DLK1. Group 6 virgins replicate group 1 virgins, and group 8 replicate group 2 since they have normal conceptus-derived plasma DLK1 expression. Group 7 females were crossed to a *Dlk1^-/-^* sire, thus lack conceptus-derived plasma DLK1 expression. Groups 3, 5 and 4 were *Dlk1^-/-^* females and so did not have a functional copy of DLK1. Group 5 females had normal conceptus-derived plasma DLK1 expression while group 4 females were crossed to a *Dlk1^-/-^* sire and so lacked conceptus-derived plasma DLK1. **(B)** To test the effect of normal pregnancy on liver and plasma lipids, we conducted three replicate comparisons between virgin and pregnant groups of matched maternal *Dlk1* genotype. **(C)** The influence of conceptus-derived circulating DLK1 on the normal pregnant lipidome was then investigated by two replicate comparisons between pregnant groups with and without a functional fetal *Dlk1* copy in matched genotype dams (“Fetal Effect”). Similarly, the influence of DLK1 derived from maternal tissues on the normal pregnant lipidome was tested by comparing DLK1+ with DLK1- dams with matched fetal DLK1 production (“Maternal Effect”).

### The maternal circulating and hepatic lipid profiles undergo broad changes to triglycerides and phospholipids in late pregnancy

We performed an untargeted lipidomics screen of plasma and liver samples from the cohort described above, utilising Direct Infusion high-resolution Mass Spectrometry (DI-MS, see Materials and Methods) and classified the resulting lipidomics dataset into major lipid classes, comparing their relative abundance between the three genotype-matched virgin and pregnant group-pairs (Supplementary Tables S1,2). As expected in the catabolic phase of pregnancy, triglycerides (TG) were considerably higher in plasma and liver (Figure 2A, B). By contrast, the most abundant lipid class, phosphatidylcholine (PC), did not show consistent changes in liver or plasma as a result of pregnancy (Supplementary Figure S1A-D). Similarly, phosphatidylethanolamine (PE), a major membrane phospholipid, was only increased in plasma, and was unchanged in the livers of pregnant mice (Figure 2C, Supplementary Figure S1E). We observed shifts in less abundant phospholipids; cardiolipins (CL), which constitute the principal lipid component of mitochondria, were reduced in the livers of pregnant groups (Figure 2D). Moreover, the *lyso-*phospholipids (LPC, LPE and LPS), which are formed by the hydrolysis of one fatty acid residue from a phospholipid, as well as plasmalogens (ether-linked lipids, e.g. PC-O), were all decreased in the plasma of pregnant mice when compared to virgin females (Figure 2E-H). The similarities in abundance between conditions for hepatic PC and PE, and the reduced hepatic CL in pregnant groups were supported by phosphorus NMR profiling of pooled liver samples (Supplementary Table S3).

**Figure 2.**
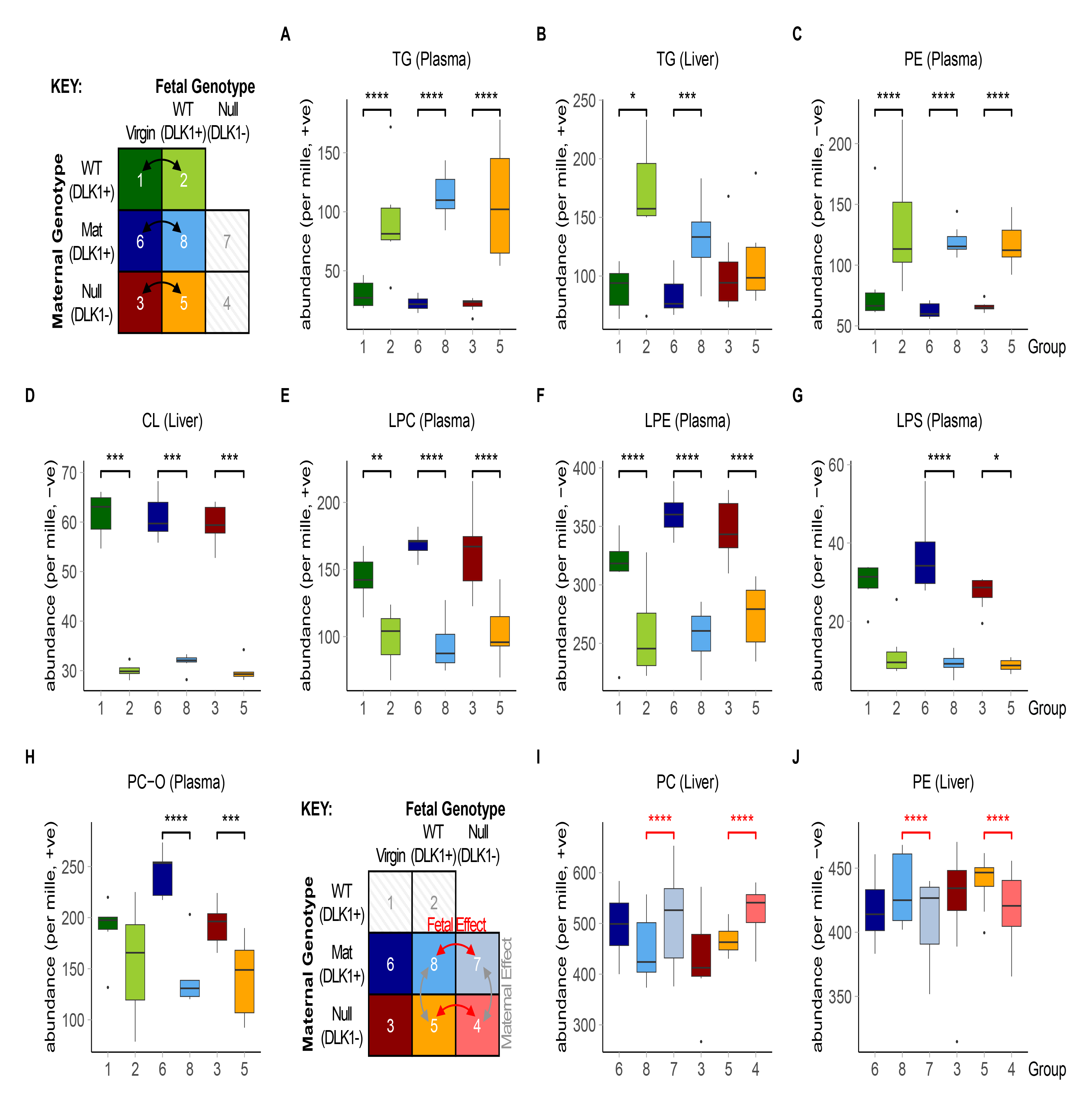
Lipid classes that change in the liver or plasma of mice in normal and *Dlk1*-manipulated pregnancy. **(A-J)** Grouped relative abundance of lipid classes that are significantly different in liver or plasma in pregnant mice (15.5 dpc) compared to virgin controls **(A-H)**, and in pregnant mice that lack fetal or maternal-derived DLK1 protein **(I-J)**. Significance was only considered if identified in at least two genotype-matched replicate group comparisons. All class data are found in Supplementary Tables S1-2. Data is presented as boxplots with whiskers showing 1.5*IQR and outliers plotted individually. Two-way ANOVA with Sidak’s multiple comparisons test was performed to determine significant class shifts between experimental groups (* p-value <0.05; ** p-value <0.01; *** p-value <0.001; **** p-value <0.0001). Statistical tests were performed independently per ionisation mode and per genotype-matched replicate comparison. n=5-8 per group. CL, cardiolipin; LPC, *lyso*-phosphatidylcholine; LPE, *lyso*- phosphatidylethanolamine; LPS, *lyso*-phosphatidylserine; PC, phosphatidylcholine; PC-O, PC plasmalogen; PE, phosphatidylethanolamine; TG, triglyceride.

Since class-wide changes in hepatic PC and PE were not evident when comparing pregnant and virgin groups, we investigated whether maternal phospholipids were altered based on their fatty acid compositions. Phospholipids contain fatty acid chains in two distinct biochemical positions on the glycerol moiety; the *sn-*1 position is typically occupied by a saturated fatty acid whereas the *sn-*2 position is more often occupied by an unsaturated residue. DI-MS is unable to distinguish which individual fatty acids occupy these positions, but rather reports the sum of carbon chains and unsaturated bonds on both positions (e.g. 40:6 in the example in Figure 3A). We grouped PC lipids by the total number of double bonds on their fatty acid chains and found that those with four or more double bonds were increased while those with three or fewer were decreased in pregnant groups (Figure 3B, Supplementary Figure S2A). Thus, we found a substantial shift in abundance between subclasses of PC. This unsaturation-specific directionality was apparent in both liver and plasma. A similar abundance shift was evident for PE lipids in liver (Figure 3C). In plasma, where PEs are much less abundant, PEs with four or more double bonds increased in abundance, however, those with three or fewer were unchanged in pregnancy. Hence, while triglycerides show a characteristic class-wide increase in the livers and circulation of pregnant mice, the two major phospholipid classes (∼80-90% of total phospholipid) instead undergo selective shifts to their fatty acid composition in favour of a more unsaturated phenotype, suggesting pregnancy-associated changes to the control of the system.

**Figure 3.**
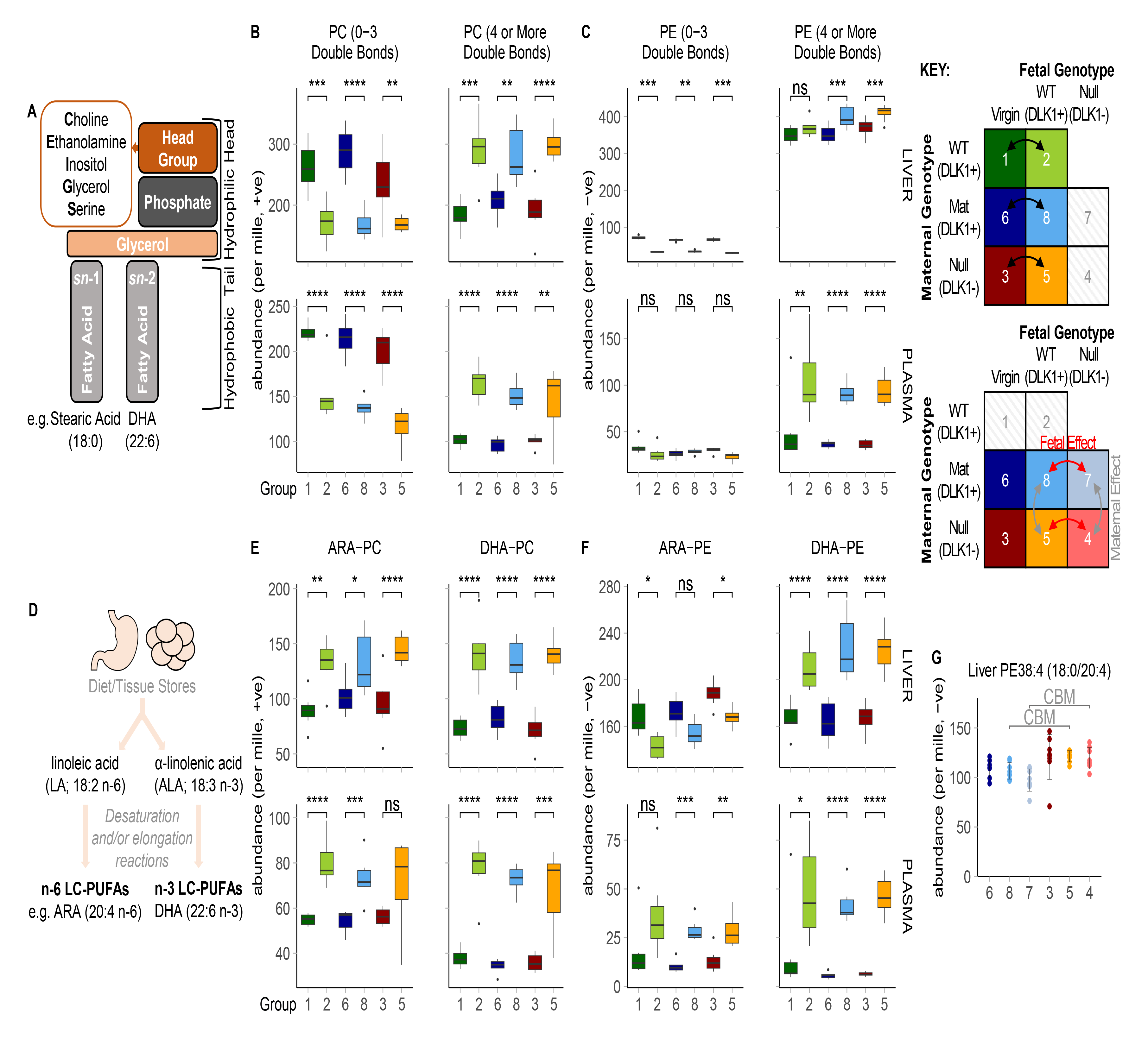
Changes to the fatty acid composition of PC and PE lipids in normal and *Dlk1*-manipulated pregnancy. **(A)** Phospholipid structure. Phospholipids contain various head groups and are commonly associated with a saturated or monounsaturated fatty acid at the *sn*-1 position and a monounsaturated or polyunsaturated fatty acid at the *sn*-2 position. **(B-C)** Grouped relative abundance of PC **(B)** and PE **(C)** lipids that contain fatty acids with a combined total of three or fewer double bonds (left) or four or more double bonds (right) in the liver and plasma of virgin and pregnant (15.5 dpc) groups. PC data in the negative ionisation mode is shown in Supplementary Figure S2A**. (D)** Schematic of n-6 and n-3 LC-PUFA synthesis from essential fatty acid precursors. **(E-F)** Grouped relative abundance of the most common PC **(E)** and PE **(F)** lipids that specifically contain ARA (left) or DHA (right), as identified by targeted LC-MS/MS analysis. List of included lipids are found in Supplementary Table S7. Grouped abundance data is presented as boxplots (whiskers showing 1.5*IQR and outliers plotted individually) and two-way ANOVA with Sidak’s multiple comparisons were performed for each genotype-matched virgin vs pregnant comparison (* p-value <0.05; ** p-value <0.01; *** p-value <0.001; **** p-value <0.0001). Significance was only considered if identified in at least two genotype-matched replicate comparisons. **(G)** Relative abundance of PE38:4 which was identified as a candidate biomarker (CBM) that distinguishes the livers of pregnant dams lacking maternal-derived DLK1 from those with normal expression of maternal-derived DLK1. CBMs are classified as lipids that passed both Bonferroni-adjusted *t*-tests (liver threshold, p = 0.00234) and sparse partial least squares discriminant analysis in two genotype-matched replicate comparisons (see Supplementary Table S5 for full list of CBMs). CBM data is shown as individual relative abundance values with ± SD error bars and *sn*-1/*sn*-2 fatty acid compositions were assigned by targeted LC-MS/MS analysis (Supplementary Table S6). All statistical tests were performed independently per ionisation mode and per genotype-matched replicate comparison. n=5-8 per group. ARA, arachidonic acid; DHA, docosahexaenoic acid; PC, phosphatidylcholine; PE, phosphatidylethanolamine.

We next evaluated the lipid class profile of pregnancy using our *Dlk1*-model of perturbed maternal lipid adaptations. While triglycerides and less abundant phospholipid classes were unchanged between DLK1+ and DLK1- pregnant groups, we observed a shift in the most abundant phospholipid classes in the maternal liver (Supplementary Table S2). Specifically, pregnant dams that lacked fetal-derived circulating DLK1 had higher amounts of PC and reduced amounts of PE in their livers, compared to those with normal circulating DLK1 (Figure 2I, J). Unlike the saturation-specific effect of normal pregnancy, the DLK1-dependent shift in abundance of these phospholipids was due to small alterations to several lipids within the class (Supplementary Figure S2B-D), suggesting that DLK1 derived from fetal tissues has a class-wide influence on maternal hepatic PC and PE abundance in pregnancy.

### Circulating ARA and DHA in late pregnancy is driven by the selective hepatic production and export of phospholipids, not triglycerides

To identify the molecular lipid species that best distinguished pregnant from virgin groups we next applied a stringent statistical pipeline, known as “Candidate Biomarker” (CBM) discovery (22). In at least two replicate virgin vs pregnant group comparisons, lipids that passed sparse Partial Least Squares Discriminant Analysis (sPLS-DA) multivariate test, followed by an FDR-corrected Student’s *t*-test, were classified as CBMs. Of the resulting 43 CBMs, 29 were PCs or PEs, suggesting that the observed shifts in maternal phospholipids are represented by select PC and PE species (Supplementary Table S4).

We used targeted LC-MS/MS to quantify the FA composition of PC and PE CBMs that were driving the unsaturation-specific shifts in maternal phospholipids (Supplementary Table S6). Notably, those that contained four or more double bonds, PC(37:4, 38:6, 39:6, 40:6) and PE(40:6), were all raised in both liver and plasma, and contained DHA or ARA (Table 1; Supplementary Figure S3A). These data reveal a selective rise in phospholipids that contain the two most biologically important LC-PUFAs in pregnancy. By contrast, PCs that contained three or fewer double bonds, PC(34:2, 35:3, 36:3), were lower in pregnant groups and contained LA (18:2 n-6), the diet-derived precursor fatty acid in n-6 LC-PUFA synthesis (Table 1; Supplementary Figure S3B; Figure 3D). The specific alterations in the profile of PUFAs in PC suggests that the hepatic LC-PUFA pathway and their incorporation into phospholipids are key regulatory targets in late pregnancy. Moreover, the parallel changes in liver and plasma implicate the hepatic transfer of these particular lipids into the circulation.

**Table 1:**
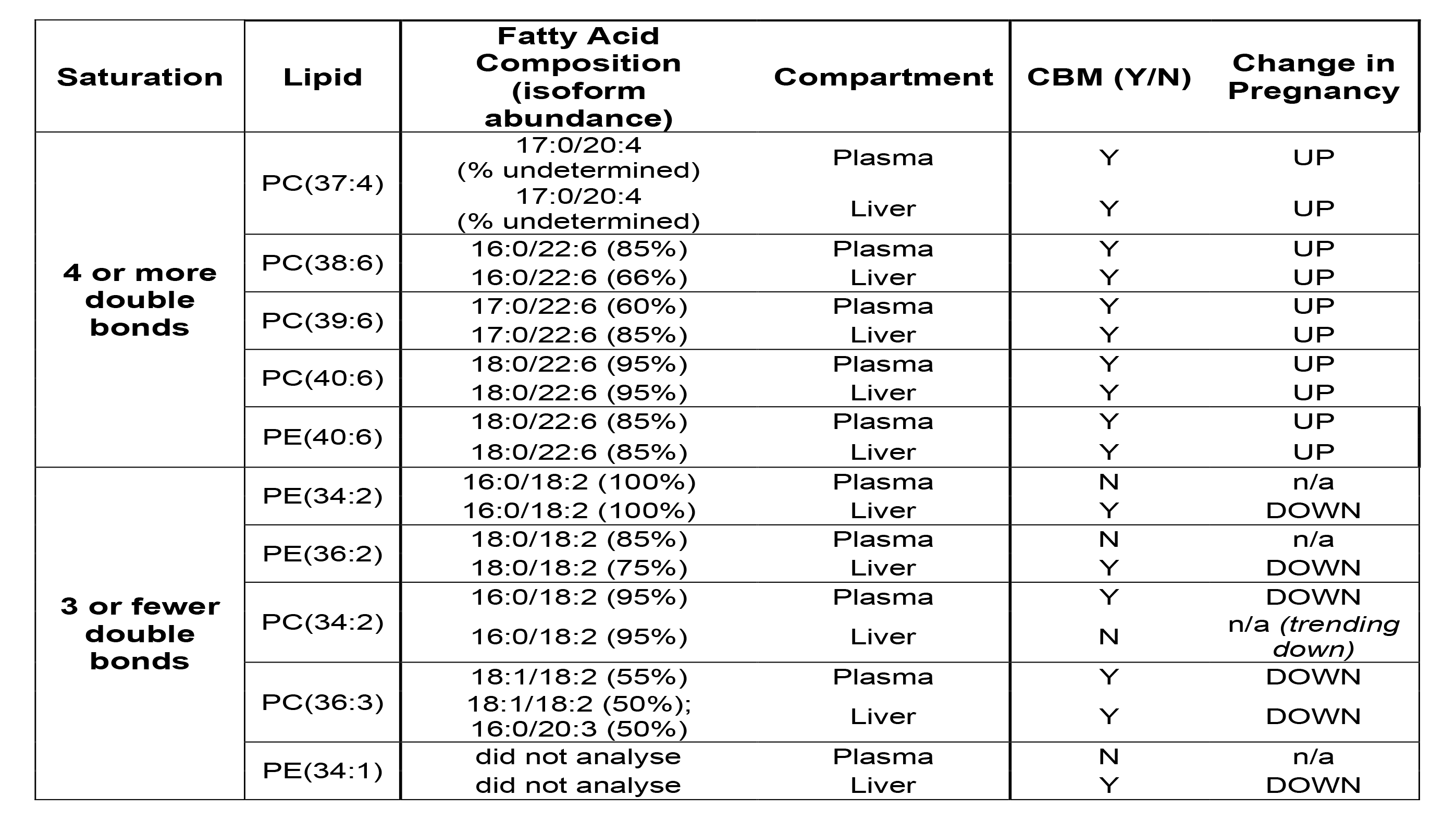
PC and PE species of interest that were identified as candidate biomarkers associated with pregnancy. PC and PE candidate biomarkers (CBMs) that distinguish pregnant groups (15.5 dpc; groups 2, 8 and 5) from virgin groups (groups 1, 6 and 3) were selected based on their level of unsaturation and the tissue compartment in which they were identified. *sn*-1/*sn*-2 fatty acid compositions were assigned using the most abundant isoform identified from targeted LC-MS/MS analysis in plasma and liver (percentage of total lipid/glyceride signal indicated in brackets; see Supplementary Table S6). CBMs were classified as lipids that passed both Bonferroni-adjusted *t*-tests (liver threshold, p = 0.00234; plasma threshold, p = 0.00283) and sparse partial least squares discriminant analysis in at least two genotype-matched virgin vs pregnant comparisons (see Supplementary Table S4 for full CBM list). Individual CBM plots are depicted in Supplementary Figure S3. All CBM tests were performed independently per ionisation mode and per genotype-matched replicate comparison. n=5-8 per group. PC, phosphatidylcholine; PE, phosphatidylethanolamine.

Pregnancy-associated CBMs were evaluated further by comparison to the most abundant ARA or DHA-containing PCs and PEs of the mouse liver using a published lipidomics profile (19) and targeted LC-MS/MS to identify and quantify the FAs they comprised (list of included lipids in Supplementary Table S7). When the combined abundances were compared between virgin and pregnant groups, ARA-PC, DHA-PC and DHA-PE were considerably more abundant in both maternal liver and plasma, the PC lipids showing concurrence between the two compartments (Figure 3E, F). These data highlight the wide-spread increase of LC-PUFA-containing phospholipids in late pregnancy, and suggest that the maternal liver selectively promotes the generation and export of phospholipids that contain ARA and DHA.

Although triglycerides are considered a major supply route for LC-PUFAs to the fetus, only one minor triglyceride species, TG(56:6), was identified as a CBM that was different between virgin and pregnant groups (Supplementary Table S4). Hence, in contrast to phospholipids, the observed class-wide rise in plasma and hepatic triglycerides is interpreted as the sum of minor increases by each individual triglyceride. As triglycerides and phospholipids have very different ionisation efficiencies and it is therefore difficult to compare concurrent changes in both groups, it is sometimes mores useful to determine within group changes. Indeed, when we split triglycerides into those that do and those that do not contain LC-PUFAs, both types were raised in pregnant groups compared to virgins (Table 2). Crucially, data in Table 2 highlights how the proportional abundance of LC-PUFAs is considerably higher in PCs compared to triglycerides and diglycerides. We therefore propose that the general rise in maternal triglycerides increases the supply of all fatty acids to the fetus, while the more regulated shifts in PC composition substantially increases the availability of LC-PUFAs for fetal uptake.

**Table 2:** Relative abundance of PC and triglycerides that contain a LC-PUFA in the liver and plasma of virgin and pregnant mice. The relative abundance of PC, triglycerides and diglycerides (which represent fragmented triglycerides) were grouped based on whether they contain a LC-PUFA. LC-PUFA-containing PCs were assigned based on targeted LC-MS/MS analysis (Supplementary Table S6). Triglyceride signals with 54 or more carbons and 5 or more double-bonds, and diglyceride signals with 38 or more carbons and 5 or more double-bonds, were expected to contain a LC-PUFA. Data is presented in the positive ionisation mode to highlight the proportion of LC-PUFAs in triglycerides compared to PC. Data is combined for pregnant (15.5 dpc; groups 2, 8 and 5) and virgin (groups 1, 6 and 3) groups and presented as mean relative abundance (±SD). n=5-8 per group. DG, diglyceride; LC-PUFA, long-chain polyunsaturated fatty acid; PC, phosphatidylcholine; PE, phosphatidylethanolamine; TG, triglyceride.

| Compartment | Class | Relative Abundance (mean ± sd, per mille) | | | |
| --- | --- | --- | --- | --- | --- |
|  |  | LC-PUFA-containing |  | Non-LC-PUFA-containing |  |
|  |  | VIRGIN | PREGNANT | VIRGIN | PREGNANT |
| Plasma | PC | 99.66 (6.93) | 154.13 (25.94) | 211.09 (19.65) | 135.9 (25.65) |
|  | TG | 1.19 (1.4) | 24.21 (9.5) | 23.33 (7.99) | 79.8 (29.81) |
|  | DG | 1.46 (3.4) | 7.19 (2.74) | 133.79 (37.22) | 183.6 (51.03) |
| Liver | PC | 194.52 (31.5) | 290.5 (39.93) | 262.68 (46.82) | 169.41 (21.71) |
|  | TG | 11.5 (5.04) | 19.39 (6.28) | 80.37 (21.6) | 112.6 (38.13) |
|  | DG | 33.25 (5.96) | 38.93 (8.19) | 208.79 (33.19) | 201.5 (32.53) |

### The n-6 PUFA composition of PEs undergo liver-specific modifications in late pregnancy

An exception to the rise in hepatic and circulating LC-PUFA-phospholipids in late pregnancy was observed for ARA-containing PE, which was increased in plasma but reduced in livers of pregnant mice (Figure 3F). We tested whether the shifts in individual PE species were specific to one of the two maternal compartments. With the exception of the highly abundant DHA-containing lipid, PE(40:6), which increased in both liver and plasma, all other PE CBMs were only identified in one compartment (Supplementary Table S4). This is in direct contrast to the parallel changes to nearly all PC CBMs.

Two abundant PE lipids that contain LA (18:2 n-6), PE(34:2, 36:2), were lower in pregnant groups, but unlike LA-containing PC CBMs, this fall was specific to the maternal liver (Table 1, Supplementary Figure S3B). Of the three most abundant ARA-containing PE species, PE(38:5) was lower only in the livers of pregnant mice, while PE(38:4, 36:4), were higher, but only in plasma. Of note, around 20% of PE(38:5) contains EPA (FA(20:5 n-3)), rather than ARA, as determined on the basis of fragmentation data obtained by targeted LC-MS/MS (Supplementary Table S6). These data together suggest a species-specific regulation of ARA-PE and EPA-PE in the liver, and warrants further investigation into isoform-specific adaptations to hepatic PE in pregnancy.

When we used the CBM analysis to compare individual lipids between DLK1+ and DLK1- pregnant groups, PE(38:4) was identified as a CBM that was increased in the livers of pregnant mice that lacked DLK1 in maternal tissues (Figure 3G, Supplementary Table S5). This lipid is notably the most abundant of all isoforms of PE, and thus changes in it represent not only a clear shift in the configuration of the system but also a sizable proportion of hepatic ARA. Taken together, these data reveal the complex control of n-6 PUFAs in PE in the maternal liver, which may in part be influenced by maternally-produced DLK1.

### ARA-containing phospholipids are hydroxylated in the maternal liver to generate bioactive lipids of the cyclooxygenase pathway

The enzymatic cleavage of PUFAs, principally ARA, from membrane phospholipids is the rate-limiting step in in the generation of bioactive PUFA metabolites (23). To investigate whether the maternal liver uses ARA-phospholipids to enhance their biosynthesis, a PUFA metabolite panel was run on the virgin and pregnant liver samples. LC-MS/MS was used to measure the concentrations of 38 metabolites that are synthesised from PUFAs via cyclooxygenase (COX), lipoxygenase (LOX), Cytochrome P450 (CYP450) or non-enzymatic beta-oxidation pathways (Figure 4A). To identify metabolites that best distinguished pregnant from virgin groups, we applied the CBM discovery statistical technique, which identified four COX-synthesised metabolites (prostaglandin D2, prostaglandin E2, prostaglandin F2α and thromboxane B2) and two LOX-synthesised metabolites (5- hydroxyeicosatetraenoic acid and lipoxin A4) as CBMs, all of which were higher in the maternal liver (Figure 4B-H). Immunohistochemistry on sections from virgin and pregnant livers confirmed the hepatic expression of the COX-1 protein, which was located within and in close proximity to endothelial cells (Figure 4I, J). Notably, all six CBMs were derived from ARA, which mirrors the generally higher levels of ARA-derived metabolites observed in the COX, LOX and CYP450 pathways (Figure 4A). Despite the influence of DLK1 on hepatic PE(18:0/20:4), no effect was observed on PUFA metabolite concentration. Taken together, the reduced hepatic abundance of ARA-containing PE in pregnancy may be a result of increased PUFA metabolite biosynthesis in the late pregnant liver, particularly within the cyclooxygenase pathway.

**Figure 4.**
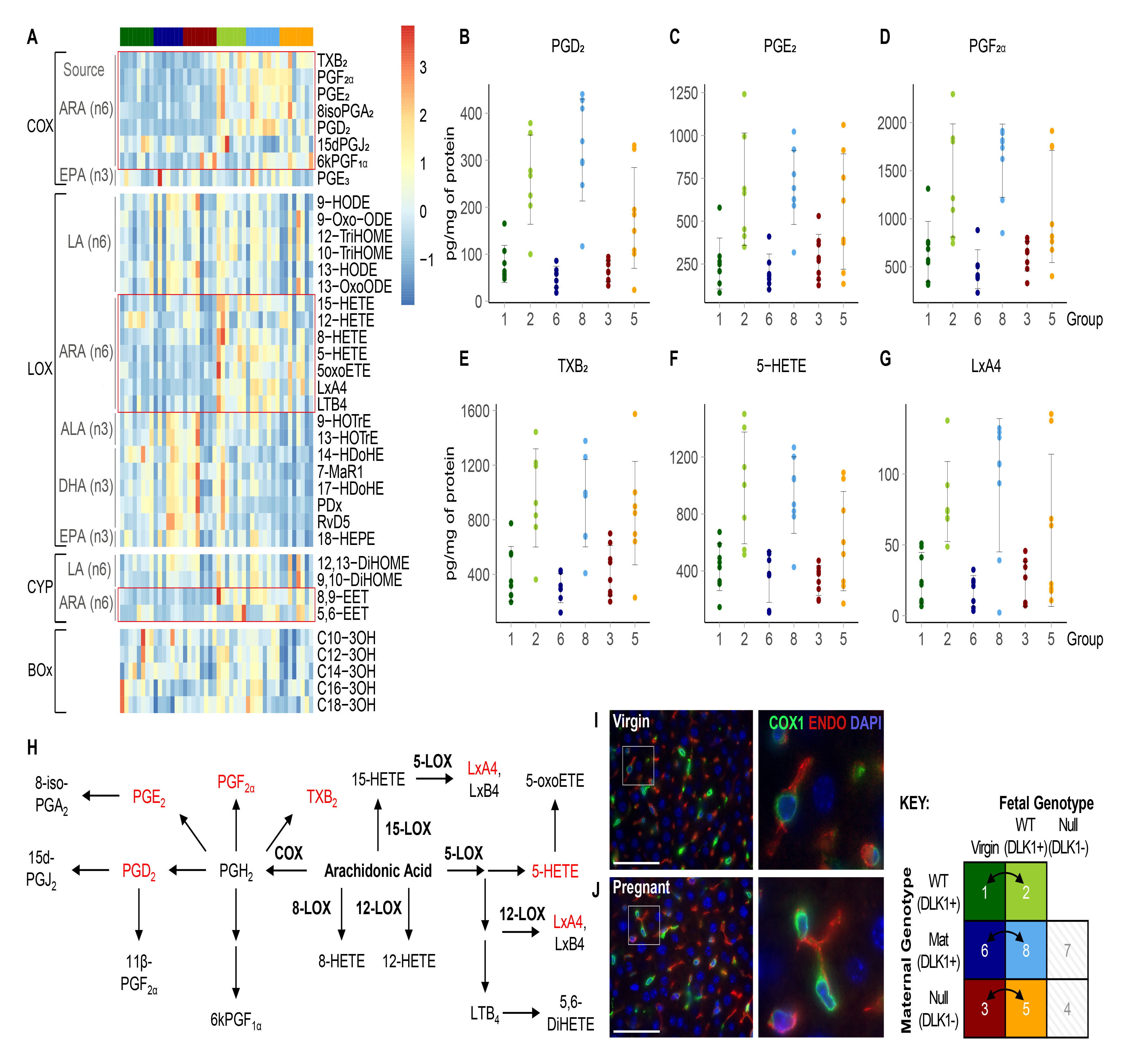
Hepatic profile of PUFA metabolites in virgin and pregnant mice. **(A)** Heatmap showing all PUFA metabolite concentrations measured in the livers of virgin and pregnant (15.5 dpc) groups. Rows are organised by synthesis pathway and further sub-divided by the precursor PUFA from which the metabolite is sourced. Red boxes indicate ARA-derived metabolites. Values are z-transformed across samples. **(B-G)** PUFA metabolites identified as candidate biomarkers (CBMs) that distinguish pregnant from virgin livers. CBMs are classified as metabolites that passed both Bonferroni-adjusted *t*-tests (p-value threshold = 0.00811) and sparse partial least squares discriminant analysis. CBM data is shown as individual concentration values with ± SD error bars. CBM tests were performed individually per genotype-matched virgin vs pregnant comparison. **(H)** Schematic tree diagram of all measured metabolites in the LOX and COX pathways that are derived from ARA. Metabolites identified as CBMs of pregnancy are indicated in red. **(I-J)** Immunohistochemistry showing COX1 and endomucin co-expression in an independent cohort of virgin **(I)** and pregnant **(J)** livers. Scale bars represent 500 μm for low-magnification images and 50 μm for enlarged images. All metabolites are defined in the Materials and Methods. ALA, α-Linolenic acid; ARA, arachidonic acid; BOx, beta-oxidation; COX, cyclooxygenase; CYP, Cytochrome P450; DHA, docosahexaenoic acid; ENDO, endomucin; EPA, eicosapentaenoic acid; LA, linoleic acid; LxA4, lipoxin A4; LOX, lipoxygenase; PGD_2_, prostaglandin D_2_; PGE_2_, prostaglandin E_2_; PGF_2α_, prostaglandin F_2α_; TxB2, thromboxane B2; 5-HETE, 5-hydroxyeicosatetraenoic acid.

### Hepatic biosynthesis and export of LC-PUFA-phospholipids is transcriptionally upregulated in pregnancy

We speculated that enzymatic pathways involved in the synthesis and export of LC-PUFA-phospholipids were crucial targets for adaptation of the late gestational liver. To investigate candidate pathways, we re-analysed a microarray dataset that compared liver transcriptomes between virgin and pregnant mice at 14.5 dpc (24) (Figure 5A), which resulted in 3272 differentially expressed genes (Figure 3B, C; Supplementary Table S8). Using a targeted approach, the dataset was assessed for transcriptional changes to rate-limiting steps within candidate pathways of interest, as depicted in Figure 5D (Supplementary Table S9). Target genes were then validated by real-time quantitative PCR (RT-qPCR) analysis in virgin and pregnant liver samples from the current study (Figure 5E-I).

**Figure 5.**
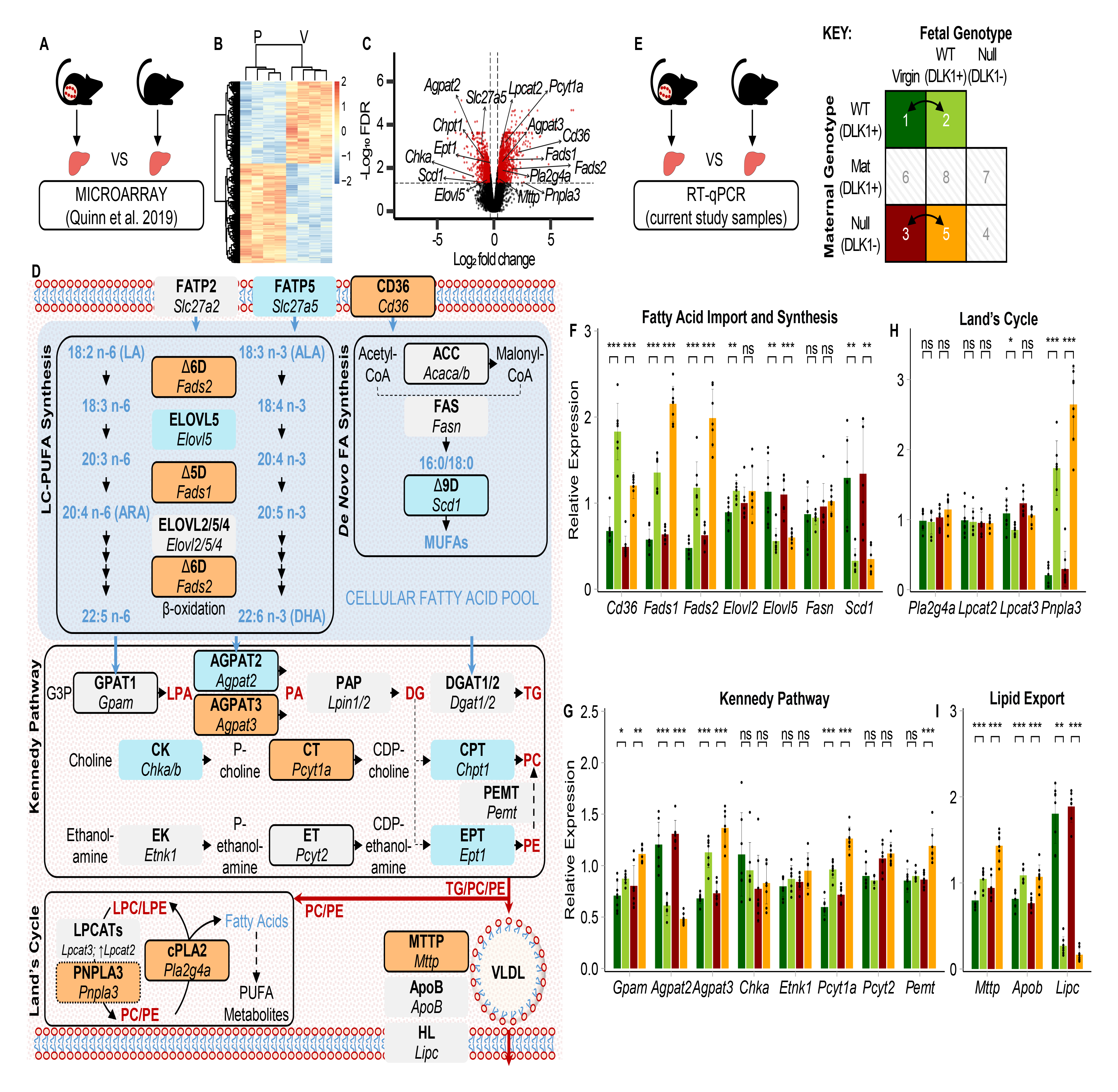
Transcriptional analysis of LC-PUFA-phospholipid biosynthetic pathways in the pregnant liver. **(A)** Schematic depicting the re-analysis of a published whole-transcriptome microarray dataset of livers from pregnant (14.5 dpc) and virgin mice (n=4/group). **(B)** Heatmap showing z-transformed values of 3272 differentially expressed genes (DEGs) based on a p-value (corrected for multiple hypothesis testing based on the Benjamini–Hochberg procedure) threshold of 0.05 and a fold change threshold of 1.25. List of DEGs are found in Supplementary Table S8. **(C)** Volcano plot showing DEGs (red) annotated with biosynthetic genes of interest. **(D)** Diagrammatic summary of hepatic lipid biosynthetic pathways chosen for targeted transcriptomics analysis annotated with microarray expression data (orange box = DEG that increased in pregnant livers; blue box = DEG that decreased in pregnant livers; grey box = gene not identified as differentially expressed). Known rate-limiting steps are indicated by thick black borders. Gene symbols are defined in Supplementary Table S9. **(E)** Schematic depicting real-time quantitative PCR (RT-qPCR) validation of candidate gene expression in livers from two genotype-matched virgin vs pregnant group comparisons (n=7- 8/group). **(F-I)** RT-qPCR data is split into fatty acid import and synthetic pathway genes **(F)**, Kennedy pathway genes **(G)**, Land’s Cycle genes **(H)** and lipid export pathway genes **(I)**. RT-qPCR data was normalised to housekeeping gene expression (*Tuba1*, *Tbp* and *Hprt*) and is shown as mean relative expression ± SD. Groups were called significantly different by Mann-Whitney U test (* p-value <0.05; ** p-value <0.01; *** p-value <0.001). DG, diglyceride; LPA, *lyso*-phosphatidic acid; LPC, *lyso*-phosphatidylcholine; LPE, *lyso*- phosphatidylethanolamine; PA, phosphatidic acid; PC, phosphatidylcholine; PE, phosphatidylethanolamine; TG, triglyceride.

We first investigated the source of hepatic LC-PUFAs in pregnancy. ARA and DHA may be directly imported into the liver from the diet or adipose tissue mobilisation, or they may be synthesised from n-3 and n-6 PUFA precursors using the LC-PUFA synthesis pathway (9). For either case, the hepatic ARA and DHA pool requires FA influx, and we observed a two-fold increase in maternal gene expression of CD36, the major regulatory transporter of hepatic fatty acid uptake (Figure 5D, F). Gene expression of fatty acid transfer proteins were unchanged (*Slc27a2*) or decreased (*Slc27a5*) in the microarray dataset (Figure 5D). Crucially, *Fads1* and *Fads2* were upregulated in the maternal liver, which encode the rate-limiting delta-5 and delta-6 desaturases in LC-PUFA synthesis (Figure 5D, F). LC-PUFA elongation enzymes were either unchanged (*Elovl2*), or down-regulated (*Elovl5*) (Figure 5D, F). The rate-limiting desaturase in monounsaturated fatty acid formation, *Scd1,* was considerably down-regulated in pregnancy, while no other changes were observed in the *de novo* fatty acid synthetic pathway (Figure 5D, F), consistent with the lipidomics data that shows a selective increase in LC-PUFAs. These findings together support the influx and subsequent desaturation of n-3 and n-6 PUFA precursors in the maternal liver.

The hepatic pool of ARA and DHA are incorporated into lipids through synthetic pathways. The Kennedy Pathway describes the *de novo* synthesis of both triglycerides and phospholipids, wherein phosphatidic acid (PA) is generated as a common precursor (25). The initial committing step in this pathway was upregulated in the pregnant liver (*Gpam;* Figure 5G). For the second PA-generating step, the two genes for major hepatic PA-acyltransferases were oppositely regulated by pregnancy. *Agpat2,* encoding LPA-acyltransferase 2 was downregulated, whereas *Agpat3* was induced (Figure 5D, G). Importantly AGPAT2 and AGPAT3 have been reported to incorporate different fatty acids into PA, with AGPAT3 shown to selectively generate DHA-PA (26). Diglycerides generated from PA are then used in the 3-step PC and PE synthetic pathways. No difference was observed for the initial committing steps (*Chka*, *Etnk1;* Figure 5G), but the second rate-limiting step in PC synthesis was increased in the pregnant liver (*Pcyt1a;* Figure 5D, G). The equivalent rate-limiting step in PE synthesis was unchanged (*Pcyt2;* Figure 5D, G). Thus, selective adaptations to the Kennedy Pathway may promote AGPAT3-mediated synthesis of DHA-containing PCs in pregnancy.

While incorporation of DHA into phospholipids is thought to occur mainly as a result of the Kennedy pathway, ARA-containing phospholipids are predominantly generated by the Land’s cycle, which cleaves and re-acylates fatty acids on the *sn-2* position of fully synthesised phospholipids (25). Incorporation of ARA into phospholipids may be achieved by activation of re-acylation enzymes, of which *Lpcat3* has known specificity for ARA (27,28). However, we observed only small shifts in the expression of genes encoding known phospholipid acyltransferase enzymes (Figure 5D, H). Enzymes that cleave the *sn*-2- position of phospholipids, phospholipase A_2_ (PLA_2_) lipases, are members of a large, complex and not fully characterised protein family (29,30). Research into PLA_2_ function has largely focused on their role in liberating ARA for PUFA metabolite synthesis, for which the PLA_2_ isoform with specificity to ARA in the *sn*-2 position, encoded by *Pla2g4a*, catalyses the known rate limiting step. Elevated expression of *Pla2g4a* in the pregnant liver (Figure 5D) is consistent with the observed pregnancy-specific rise in ARA-derived metabolites (Figure 4A). PLA_2_ isoforms involved in cleaving other *sn*-2 fatty acids from phospholipids for subsequent ARA incorporation are not established. Two other PLA_2_-encoding genes, *Pla2g7* and *Pla2g15* were upregulated in the pregnant liver (Supplementary Table S9), yet their substrate specificities and roles have not been fully characterised. Therefore, we were unable to uncover any evidence for selective increase in the Land’s cycle activity that favour increased ARA incorporation in phospholipid of the pregnant liver, at the transcriptional level.

We additionally considered alternative phospholipid synthetic pathways, including the liver-specific conversion of PE into PC via phosphatidylethanolamine methyl transferase (PEMT). Both gene and protein expression were unchanged in pregnant livers (Figure 5D, G; Supplementary Figure S4A, B). Of note, *Pnpla3* was ∼5-fold upregulated in the pregnant liver (Figure 5D, H). This gene encodes a fatty acid remodelling enzyme which has recently been shown to transfer LC-PUFAs (including DHA and ARA) from liver triglycerides to the phospholipid compartment via alternative re-acylation pathways (31).

Ultimately, the parallel rise in triglycerides and LC-PUFA-phospholipids, particularly PC, between liver and plasma supports the lipoprotein-mediated export of these lipids. Gene expression of VLDL-specific proteins, ApoB and MTTP, were increased in the maternal liver, while that of hepatic lipase (*Lipc*), which hydrolyses circulating VLDL, was decreased (Figure 5D, I). Taken together, our targeted transcriptional analysis demonstrates the regulated adaptations to maternal hepatic lipid pathways that promote PUFA influx, PUFA-specific desaturation, *de novo* biosynthesis of LC-PUFA-containing PCs and lipid export pathways.

### LC-PUFA-phospholipid synthesis is transcriptionally co-regulated by the liver X receptor in late pregnancy

In our RT-qPCR analysis that compared two genotype-matched virgin vs pregnant livers, one pregnant group lacked DLK1 derived from maternal tissues (group 5; Figure 5E-I). We noticed that for many LC-PUFA-phospholipid biosynthetic genes that were altered by pregnancy, the effect was greater in this group compared to the wild-type pregnant mice (group 2). Considering the observed effect of DLK1 on PC and PE class levels (Figure 2I, J), and on the ARA-PE species (PE(18:0/20:4); Figure 3G), we investigated whether DLK1 affects the transcriptional adaptations promoting LC-PUFA-phospholipid synthesis in the late gestational liver.

Pregnant mice lacking maternal-derived DLK1 exhibited increased gene expression of both rate-limiting PUFA desaturase enzymes, delta-5 and delta-6 desaturase (*Fads1, Fads2;* Figure 6A), the rate-limiting step in PC biosynthesis (*Pcyt1a;* Figure 6B), the proposed TG-phospholipid trans-acylase (*Pnpla3*, Figure 6C) and the rate-limiting MTTP protein required for lipoprotein assembly (*Mttp;* Figure 6D). Alongside trends in other Kennedy pathway genes including increased *Gpam* and reduced *Agpat2* expression, this DLK1 effect notably augments the transcriptional response of normal pregnancy. We further investigated this coordinated gene response by running a multiple correlation analysis of these selected genes in pregnant and virgin mice (Figure 6E-F). The significant and similar correlations between LC-PUFA-phospholipid biosynthetic genes in pregnant mice, compared to that of virgins, suggests this pathway is a transcriptionally co-regulated process in pregnancy that can be modulated by the maternal DLK1 genotype.

**Figure 6.**
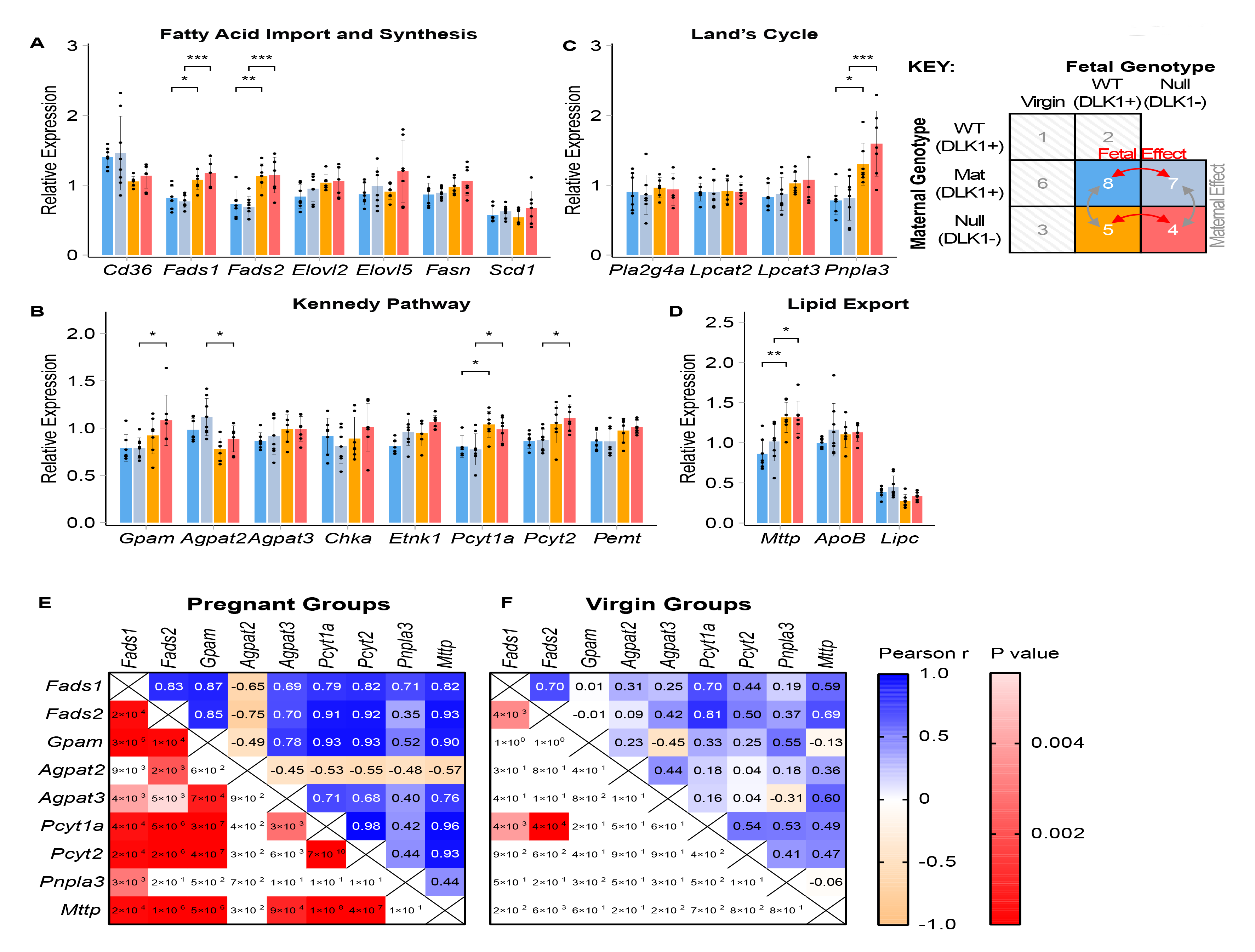
Coordinated hepatic expression of LC-PUFA-phospholipid biosynthetic genes in response to *Dlk1*-manipulation in pregnancy. **(A-D)** Real-time quantitative PCR (RT-qPCR) analysis of LC-PUFA-phospholipid biosynthetic genes in pregnant mice (15.5 dpc) in response to *Dlk1* manipulation. Genotype-matched replicate comparisons were performed to assess the effect of maternal-derived DLK1 protein or fetal-derived DLK1 protein on the expression of fatty acid import and synthetic pathway genes **(A)**, Kennedy pathway genes **(B)**, Land’s Cycle genes **(C)** and lipid export pathway genes **(D)**. Gene symbols are defined in Supplementary Table S9. RT-qPCR data was normalised to housekeeping gene expression (*Tuba1* and *Hprt*) and is shown as mean relative expression ± SD (n=7-8/group). Gene expression was compared between all four experimental groups by one-way ANOVA with Tukey’s multiple-comparison post-hoc test (* p-value <0.05; ** p-value <0.01; *** p-value <0.001). **(E-F)** Multiple Pearson’s correlations between DLK1-responsive genes in pregnant (groups 2 and 5; **E**) and virgin (groups 1 and 3; **F**) livers. P-values in shaded red boxes indicate statistically significant correlations (Bonferroni-adjusted p-value threshold = 0.0045).

To explore potential upstream regulators involved in the coordinated LC-PUFA-phospholipid transcriptional response, we assessed the differentially expressed genes derived from the virgin and pregnant hepatic microarray re-analysis for enrichment of transcription factor targets using ChIP enrichment analysis (ChEA) (32). The upregulated gene list from the maternal hepatic transcriptome was significantly enriched for 10 transcription factor pathways (Table 3, Supplementary Table S10), of which only the Liver X receptor (LXR), retinoid X receptor (RXR) and Estrogen Receptor 1 (ESR1) pathways induced genes involved in the coordinated maternal LC-PUFA-phospholipid response. Importantly, 6 key LC-PUFA-phospholipid genes (*Cd36*, *Fads1*, *Fads2*, *Pcyt1a*, *Pnpla3*, *Mttp*) were identified as targets of the master lipid regulator, LXR. Increased gene expression of the inducible LXRa isoform (*Nr1h3*) was confirmed in pregnant livers in both the re-analysed microarray dataset and in current study samples (Supplementary Table S9; Supplementary Figure 5A). However, classical hepatic LXR targets involved in reverse cholesterol transport (*Cyp7a1*, *Abcg5*, *Abcg8*, *Abca1*, *Pltp*, *Lpl*), and *de novo* lipogenesis (*Srebf1*, *Fasn*, *Scd1*) were unchanged or downregulated in pregnancy (Supplementary Table S9; Supplementary Figure 5B, C).

**Table 3.** Transcription factor pathways enriched for genes that are upregulated in the pregnant liver. Overlap analysis between transcription factor targets and upregulated differentially expressed genes (DEGs) generated from the re-analysis of a published whole-transcriptome microarray dataset of livers from pregnant (14.5 dpc) and virgin mice, using ChIP enrichment analysis (ChEA) through the Enrichr platform. Enriched DEGs were then searched for LC-PUFA-containing-phospholipid biosynthetic genes of interest. See Supplementary Table S10 for the full list of enriched DEGs in each transcription factor pathway.

| Transcription Factor Pathway | Adjusted P-Value | #Upregulated DEGs (/#All Targets) | LC-PUFA-Phospholipid Pathway DEGs |  |
| --- | --- | --- | --- | --- |
|  |  |  | # | Gene Names |
| FOXM1 | 1.05E-11 | 56/267 | 0 |  |
| IRF8 | 1.04E-10 | 217/2000 | 0 |  |
| SMRT | 2.45E-05 | 193/2000 | 0 |  |
| RXR | 7.03E-05 | 190/2000 | 3 | *Fads1, Fads2, Pnpla3* |
| LXR | 1.33E-04 | 188/2000 | 6 | *Cd36, Fads1, Fads2, Pcyt1a Pnpla3, Mttp* |
| MECOM | 2.76E-04 | 182/1951 | 0 |  |
| NCOR | 4.94E-04 | 184/2000 | 0 |  |
| CLOCK | 1.71E-02 | 47/407 | 0 |  |
| ESR1 | 1.78E-02 | 50/444 | 1 | *Fads2* |
| IRF8 | 2.91E-02 | 38/319 | 0 |  |

Reduced gene expression of classical lipogenic and cholesterol LXR targets in late pregnancy was previously described by Nikolova and colleagues (33), where both wild type (WT) and *Lxrab*^-/-^ double knockout (DKO) mutant dams showed this late-pregnancy associated reduction in LXR target transcription. Using liver samples from the Nikolova cohort (33), we quantified the expression of the six putative LXR-induced LC-PUFA-phospholipid genes at early and late gestational timepoints (Figure 7B-G). LXR DKO mice demonstrated a blunted late-gestational rise in *Fads1*, *Fads2*, *Pnpla3* and *Cd36* expression, with a non-significant trend also evident for *Pcyt1a.* Measurement of other key LC-PUFA-phospholipid genes revealed a similar attenuated late-pregnancy increase in *Agpat3* expression (Figure 7H) but no change to the fall in *Agpat2* expression (Supplementary Figure S5D). Reduced expression of *Mttp*, *Lpcat3* and *Gpam* in LXR DKO mice at all timepoints suggests them to be pregnancy-independent LXR targets (Figure 7G; Supplementary Figure S5E, F).

**Figure 7.**
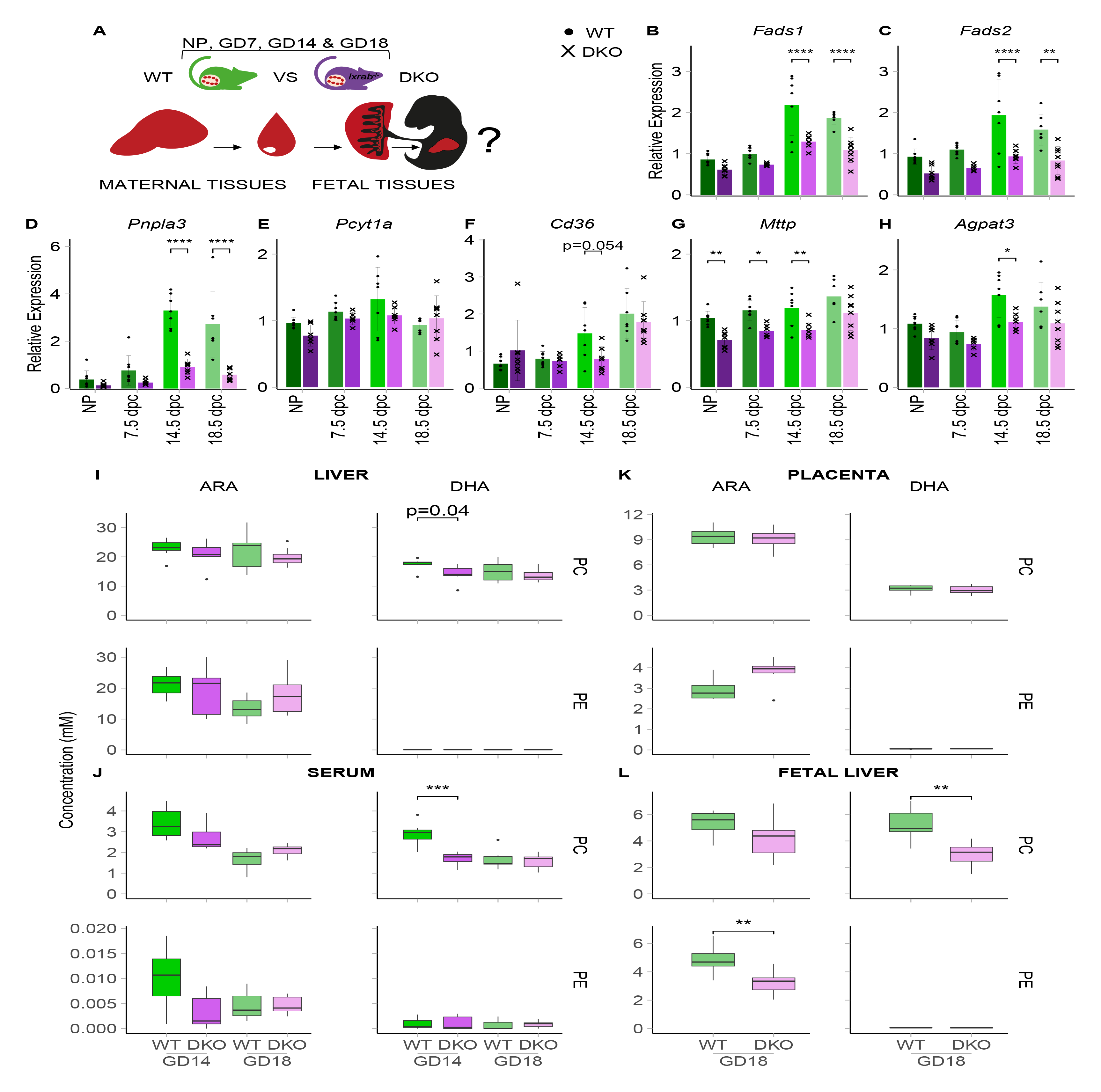
LXR-mediated activation of hepatic LC-PUFA-phospholipid biosynthesis, export and fetal transfer in late pregnancy. **(A)** Schematic of the tissues analysed from an independent cohort of wild-type (WT) and *Lxrab*^−/−^ (LXR double knockout (DKO)) mice at non-pregnant (NP) and various gestational timepoints (7.5, 14.5 and 18.5 dpc). Arrows indicate the hypothesized transfer of LC-PUFA-containing phospholipids between maternal and fetal compartments. **(B-H)** RT-qPCR analysis of hepatic LC-PUFA-phospholipid biosynthetic genes in WT and LXR DKO groups (n=6-8/group). RT-qPCR data was normalised to housekeeping gene expression (*Tuba1*, *Tbp* and *Hprt*) and is shown as mean relative expression ± SD and were compared by two-way ANOVA with Šídák’s multiple comparison (* p-value <0.05; ** p-value <0.01; *** p-value <0.001; **** p-value <0.0001). **(I-L)** Grouped concentrations of ARA-containing PC, DHA-containing PC, ARA-containing PE and DHA-containing PE lipid species selected for targeted LC-MS/MS analysis in liver **(I)**, serum **(J)**, placenta **(K)** and fetal livers **(L)** from WT and LXR DKO groups at late-gestational timepoints. Individual lipid concentrations are found in Supplementary Table S11. Grouped lipid concentrations are presented as boxplots (whiskers showing 1.5*IQR and outliers plotted individually) and Bonferroni-adjusted *t*-tests (p-value threshold = 0.025) were performed for each WT vs DKO comparison (* p-value <0.05; ** p-value <0.01; *** p-value <0.001; **** p-value <0.0001). ARA, arachidonic acid; DHA, docosahexaenoic acid; PC, phosphatidylcholine; PE, phosphatidylethanolamine.

To test whether LXR-manipulation ultimately influences LC-PUFA-phospholipid levels in pregnancy, we performed targeted LC-MS/MS in samples from the Nikolova cohort at late-gestational timepoints, specifically measuring the concentrations of the ARA and DHA-containing PC and PE lipids (Supplementary Table S11). At 14.5 dpc, LXR DKO mice have a lower concentration of DHA-PC lipids in serum alongside a non-significant fall in the liver (Figure 7I, J). A non-significant trend was also evident for ARA-phospholipids in serum. No shift was observed for these lipids at the pre-parturition 18.5 dpc timepoint, which in agreement with the observed maximal expression of LC-PUFA-phospholipid synthetic genes at 14.5 dpc compared to 18.5 dpc (Figures 7B-H), suggests that the LC-PUFA-phospholipid pathway is most active at the earlier peak catabolic phase of gestation.

Finally, we hypothesised that LXR-activation of the LC-PUFA-phospholipid transcriptional program in the maternal liver is a critical adaptive pathway that facilitates LC-PUFA delivery to the fetus (Figure 7A). We tested this hypothesis by performing targeted LC-MS/MS in fetal tissues from WT and LXR DKO dams at 18.5 dpc. While no significant changes were observed in placentas from DKO dams compared to those from WT mice (Figure 7K), DHA-PC concentrations were lower in fetal livers from DKO dams (Figure 7L). At the species level, the most abundant DHA-containing PC lipid, PC(38:6), which accounts for over 50% of all DHA-PC, was reduced in both maternal liver and serum at 14.5 dpc, and in the 18.5 dpc fetal liver (Supplementary Table S11). We also observed lower concentrations of ARA-PE in the fetal liver from DKO dams, with a non-significant fall also evident for ARA-PC. DHA-PE lipid concentrations remained unchanged across all tissues in LXR DKO mice. Our study ultimately defines an LXR-mediated pathway that selectively enriches LC-PUFAs into phospholipids to transfer ARA and DHA to the fetal compartment in late pregnancy.

## DISCUSSION

### Hepatic phospholipids, not triglycerides, are the major supplier of LC-PUFAs to the fetus

Our findings challenge the long-established role of triglycerides as the major lipid fraction that supplies LC-PUFAs to the fetus (7,34). Recent work has highlighted that phospholipids as well as triglycerides may contribute to the maternal supply of fetal LC-PUFAs; for example, administration of isotope-labelled fatty acids to pregnant women prior to cesarean section revealed that PUFAs (LA and DHA) were found in both the triglyceride and phospholipid fractions (35). Studies on rat dams in late gestation show that hepatic and serum phospholipids contain a higher proportion of LC-PUFAs compared to triglycerides (20,21). These findings together build a more complex picture of the maternal lipidome, where raised triglycerides increase the availability of all fatty acids for the fetus, while ARA and DHA are selectively channeled into phospholipids to meet the profound LC-PUFA requirements of late pregnancy. This is important because it suggests that the control point of biomagnification is in the regulation of LC-PUFA phospholipid supply by the mother.

Despite the long-standing recommendation to consume DHA-rich foods during pregnancy (36), the combined supplementation evidence does not support long-term cognitive or visual benefits in the child (37–41). By contrast, positive associations have been observed between DHA levels in maternal plasma and numerous postnatal outcomes (9). Although endogenous LC-PUFA synthesis is long-thought to only minorly contribute to total mammalian ARA and DHA requirements (8), the findings by us and others support an adaptive importance for the endogenous synthetic pathway in the pregnant liver, in order to meet the LC-PUFA needs of the fetus.

We propose two key synthetic pathways that most likely underlie the raised availability of LC-PUFA-phospholipids in pregnancy. First, we report an upregulation of LC-PUFA synthesis mediated by the delta-5 and delta-6 desaturases (encoded by *Fads1* and *Fads2*), which has previously been reported in rat pregnancy (20,42,43). The critical role of both desaturases at the transcriptional level is emphasized by the numerous associations between maternal polymorphisms within the *FADS1* and *FADS2* genes with PUFA concentrations in maternal and fetal blood (44–47). Second, we propose a pregnancy-associated selective modification to the Kennedy Pathway. In addition to the increased expression of the initial committing step that generates LPA (*Gpam*), we report an isoform switch from *Agpat2* to *Agpat3* expression, which encode acyltransferases that incorporate the second fatty acid onto LPA to generate PA, the precursor molecule to triglycerides and phospholipids. AGPAT2 is a highly expressed liver enzyme with well-established function in the Kennedy Pathway. Its preferential incorporation of LA (18:2 n-6) (26) may underlie the fall in LA-containing phospholipids observed in the current study. Although less well characterised, AGPAT3 displays distinct selectivity towards DHA (48,49). Recently Hishikawa and colleagues reported reduced levels of DHA in liver phospholipids in mice with a liver-specific deletion of *Agpat3* (50). We therefore propose an adaptive mechanism in the maternal liver that targets the Kennedy Pathway to promote the synthesis of DHA-phospholipids in pregnancy.

The concurrent changes to the same PC species between liver and plasma support a particular role for PC, which has long been reported as an LC-PUFA storage lipid (51,52), as the major transport vehicle of LC-PUFAs. Further evidence of this role can be inferred from choline investigations, which like DHA, is critical for neurodevelopment and enriched in fetal cord blood (53). Isotope-tracing in pregnant women has shown that choline enrichment in the fetal compartment is principally derived from PC that was synthesized in the maternal liver (54). Notably, this study also reported that the majority of this PC was synthesized via the PEMT pathway, building on the hypothesis that PEMT, an estrogen-responsive enzyme (55) that selectively synthesizes PCs that contain LC-PUFAs (56), is the major route for choline and DHA delivery to the fetus (57). In contradiction to this theory, we report an upregulation to the rate-limiting PC synthesis step of the Kennedy pathway (*Pcyt1a*), while no changes were observed to PEMT at the gene or protein level. Therefore, we suggest that the maternal transfer of choline and DHA is driven at least in part by the Kennedy pathway, which targets *Agpat3* and *Pcyt1a* to selectively generate DHA-PC.

In pregnancy hepatic ARA was elevated in PC but reduced in PE compared to virgin livers, implicating enhanced *sn*-2 remodeling (Land’s Cycle) in the maternal liver. The best characterized acyltransferase in this pathway, LPCAT3, shows substrate specificity towards linoleic acid and ARA (28,58,59), but a transcriptional shift in *Lpcat3* was not observed. However, we report increased expression of PNPLA3 (60). A recent knockout study proposed that PNPLA3 transfers LC-PUFAs from triglycerides to phospholipids (31), supporting the emerging significance of alternative remodeling pathways to the Land’s Cycle in LC-PUFA-phospholipid synthesis (25). Ultimately, the generation of ARA-phospholipids by LPCAT3 (27,28) and potentially by PNPLA3 (61) is vital for the normal production of hepatic VLDLs. Given the aforementioned requirement for PC in the VLDL pathway, the maternal liver may selectively synthesize ARA-PC to promote VLDL-mediated lipid export into the circulation.

The observed increase in the hepatic synthesis and export of VLDLs, which consist of a triglyceride core surrounded by a PC-rich phospholipid monolayer (62), supports the VLDL-mediated transport of hepatic triglycerides and LC-PUFA-PCs in late pregnancy. The placental uptake of PUFAs from lipoproteins is further supported by the placental expression of lipoprotein lipase (LPL) and endothelial lipase (EL), yet the transport mechanism for the liberated fatty acids remains unclear (63). Importantly, while LPLs hydrolyse triglycerides, ELs exhibit phospholipase A1 activity, removing the usually saturated fatty acid from the *sn-1* position of phospholipids. Hence, to prioritize LC-PUFA and choline transfer, the placenta may instead transport the resultant LC-PUFA-containing *lyso*-phospholipid via major facilitator superfamily domain 2 protein (MFSD2A), a newly-described *lyso-*PC symporter with high affinity for LPC(22:6) (64). Genetic deletion investigations have recognized MFSD2A as a critical means of DHA supply to the brain and retina (64,65), and clinical correlations were recently reported between placental MFSD2A expression and DHA levels in cord blood (66,67). Interestingly, we observed a 2-fold reduction in *Mfsd2a* expression in pregnant livers which may suggest a complementary reduction in maternal hepatic DHA-LPC uptake (Supplementary Table S9, Supplementary Figure S4C).

### Liver-specific function of ARA-phospholipids in pregnancy

By constructing a PUFA metabolite panel of the pregnant mouse liver, we identified raised concentrations of metabolites that were specifically derived from ARA, suggesting that the targeted de-acylation of ARA is also a crucial source for the maternal PUFA metabolite pathway. Broadly divided into the ARA-derived inflammatory mediators and the n3-PUFA-derived anti-inflammatory metabolites (68), our findings further describe the characteristic inflammatory phenotype of pregnancy. We observed a particular enrichment of classical prostanoids in the COX pathway, which have diverse housekeeping and inflammatory actions. The expression of the COX-1 isoform in hepatic endothelial cells particularly points towards their hemodynamic properties in the maternal liver (69). Given the reported associations between prostanoids and pre-eclampsia (70–72), and their critical regenerative roles following hepatic injury (73), further research into maternal hepatic bioactive PUFA metabolites may elucidate underlying mechanisms that implicate liver dysfunctions to pregnancy complications.

### Selective LXR activation in the late-gestational liver

We found that dams lacking DLK1 derived from maternal tissues had modestly increased transcription of rate-limiting steps in the hepatic synthesis and export of LC-PUFA-phospholipids. Notably, in our previous study, these mice also exhibited a reduced ability to gain adipose tissue during pregnancy. Since dietary n-3 and n-6 PUFAs are preferentially stored in adipose tissue (74), this compensatory upregulation in the liver suggests maternal adipose tissue is a critical source of the longer-chained PUFAs for hepatic ARA and DHA synthesis.

The synchronized, yet selective, transcriptional response observed in our *Dlk1* model highlights the hepatic synthesis and export of LC-PUFA-phospholipids as a regulated maternal adaptation. Using several independent models of pregnancy, our study suggests this coordinated response to be mediated by LXR, an oxysterol-activated nuclear transcription factor that forms a heterodimer with retinoid X receptor (RXR) and acts as a master regulator of lipid metabolism (75). Building on the observation from Nikolova and colleagues that classical LXR-stimulated lipid pathways, reverse cholesterol transport (RCT) and *de novo* lipogenesis, are suppressed in late gestation (33), our data proposes the hepatic LC-PUFA-phospholipid transcriptional program as an alternative LXR-induced pathway that is selectively activated in late pregnancy. The significance of this adaptive pathway is strengthened by its influence on LC-PUFA concentrations observed in the fetal liver. Amniotic-injection of PUFAs in late-gestational embryonic rats has been shown to rapidly accumulate in fetal liver and brain tissue, mainly within the phospholipid compartment (76), implicating the fetal liver as a major sink for circulating PUFAs. Hence, further research is crucial to delineate the relative trafficking pathways of maternal LC-PUFAs between and within fetal tissues. Ultimately, our study proposes that LXR-induced phospholipid synthesis in the maternal liver represents a critical source of LC-PUFAs for the developing fetus.

### Limitations

This study used Direct-Infusion Mass Spectrometry (DI-MS) to generate an unbiased lipidomics profile of the pregnant mouse liver. Since this method is unable to distinguish the fatty acid moieties associated within individual lipids, assumptions were made based on their total carbon and double-bond content. We addressed this limitation using our candidate biomarker discovery pipeline and subsequent targeted profiling by liquid chromatography-mass spectrometry (LC-MS/MS), which allowed us to acquire the fatty acid compositions of hypothesis-based lipids of interest, with the additional benefit of DI-MS signal verification. The conclusions in our study are further limited by the incomplete comparability between mouse and human lipoprotein metabolism, as the plasma cholesterol fraction is represented by high density lipoproteins (HDL) in mice and low density lipoproteins (LDL) in humans (77). However, emerging evidence supports the preferential enrichment of DHA in the HDL fraction in human pregnancy, challenging the widely assumed role for VLDL in fetal fatty acid supply (78,79). Taken together, research from us and others is highlighting how the pregnant state can profoundly contradict many foundational concepts of lipid metabolism. Future work that combines lipidomics with gold-standard isotope-tracing approaches, across multiple maternal and fetal organs, will likely reveal an extensively changed metabolic system in pregnancy.

## Acknowledgements

Funding was provided by the Medical Research Council (MRC) Grants MR/L002345/1 (MC), MR/R022836/1 (MC) and MR/J001597/1 (AFS). AK and SF were funded by the BBSRC grant BB/M027252/1. CW is a NIHR Senior Investigator (NIHR200254) and was funded by the Wellcome Trust (#092993). RA was supported by a studentship from the NIHR Biomedical Research Centre at Guy’s and St Thomas’ NHS Foundation Trust and King’s College London. We gratefully acknowledge the MetaToul (Toulouse metabolomics & fluxomics facilities, www.metatoul.fr) which is part of the French National Infrastructure for Metabolomics and Fluxomics MetaboHUB. NC received funding from the ANR (agence nationale de la recherche): ANR-18-CE14-0039 and ANR-20-CE14-0011.

## Author contributions

Conceptualization, M.C., A.K., N. C; Methodology, R. A., S. F., M.C., A.K., N. C; Investigation, R. A., S. F., M. A. M. C., S. M., A. L. M.; Writing – Original Draft, R. A., S. F., M. C.; Writing – Review & Editing, R. A., S. F., A. L. M., A. C. F-S., N. C., C. W., A. K., M. C.; Funding Acquisition, M. C., A. K., A. C. F-S., N. C., C. W.; Resources, A. K., C. W., A. C. F-S.; Supervision, M. C., N. C., A. K.

## Declaration of interests

The authors declare that they have no competing interests.

## METHODS

### 1. Animal Model

#### 1.1 Mice

Unless otherwise stated, all samples were drawn from our previous study, which generated mice with manipulations to the paternally-inherited imprinted gene, *Dlk1* (4). In brief, homozygote *Dlk1* knock-outs (Dlk1^tm1Srba^; null) (80), heterozygotes lacking the non-functional maternal *Dlk1* allele (“maternal heterozygotes” Dlk1^tm1Srba/+^; Mat) and wild-type females (Dlk1^+/+^; WT) were generated. Animals were housed in a temperature and humidity-controlled room (21 °C, 55% humidity) with a 12-12 hour light-dark cycle. Mice were fed standard chow diet (RM3, Special Diets Service; fatty acid composition of the diet is shown in Supplementary Table S12) *ad libitum* and were given access to fresh tap water daily. Following weaning at postnatal 21d, mice were housed in single-sex groups (maximum of five per cage) or occasionally singly housed, except when breeding.

#### 1.2 Experimental Cohorts

Our previous study generated various cohorts of virgin and pregnant mice at 15.5 days post coitum (dpc) with manipulated expression of the DLK1 protein. The 8 cohorts used for the current study were allocated numerical identifiers (groups 1-8) as described in Figure 1A. WT, Mat and null females were crossed with wild-type males to generate pregnant mice that have normal fetal-derived circulating levels of DLK1 and normal (groups 2 and 8) or null (group 5) local DLK1 expression from maternal tissues. Mat and null females were crossed with null males to generate pregnant mice that lacked fetal-derived circulating DLK1 and had normal (group 7) or null (group 4) local DLK1 expression from maternal tissues. Age-matched WT (group 1), Mat (group 6) and null (group 3) females were used as virgin control groups. Mice underwent terminal anesthesia with ∼0.8 mg of intra-abdominal pentobarbitol (Dolethal, Vetoquinol) per gram of body weight. Terminal plasma and liver samples were flash frozen in liquid nitrogen and stored at −80°C. All animal experiments were carried out in accordance with UK Government Home Office licencing procedures.

#### 1.3 LXR Cohorts

LXR-manipulated samples were derived from a cohort of LXR WT and *Lxrab^−/−^* (LXR double knockout (DKO)) mice generated by Nikolova and colleagues in 2017 (33). In this study, non-pregnant females and pregnant mice at 7.5, 14.5, and 18.5 dpc were euthanized following a 4-hour fast. Frozen serum, liver and fetal tissue samples were stored at −80°C and used for the current study.

### 2. Lipidomics

Lipid extraction, mass spectrometry and data processing techniques were conducted using a recently described high-throughput platform (22) and is summarized here in brief.

#### 2.1 Lipid Extraction

Livers were powdered in liquid nitrogen then homogenized using GCTU solution (6M guanidinium chloride and 1.5M thiourea) and a hand-held homogenizer (TissueRuptor II, Qiagen). Liver and plasma aliquots were injected into 96-well plates (2.4 mL/well, glass-coated, Esslab Plate+™). Internal standards (150 µL, internal standard mixture in methanol, see Supplementary Table S13 (81)), DMT (500 µL, dichloromethane, methanol (3:1) and triethylammonium chloride (500mg/L)) and water (500 µL) were added to each of the wells (96-channel pipette). Following agitation (96-channel pipette) and centrifugation (3·2 k × *g*, 2 min), 20 µL of the organic solution was transferred to a 384-well plate (glass-coated, Esslab Plate+™) and dried (N_2 (g)_). The dried films were redissolved (*tert*-butylmethyl ether, 20 μL/well, and then MS-mix, 80 μL/well) and the plate was heat-sealed and run immediately.

#### 2.2 Direct-Infusion Mass Spectrometry (Untargeted Lipidomics)

All samples were infused into an Exactive Orbitrap (Thermo, Hemel Hampstead, UK), using a TriVersa NanoMate (Advion, Ithaca US) and ionized in the positive mode at 1.2 kV for 72 seconds, followed by the negative ionisation mode at −1.5kV for 66 seconds. The analysis was then stopped and the tip was discarded between samples. Samples were run in row order and kept at 15 °C throughout acquisition. The instrument was operated in full-scan mode from *m/z* 150 to 1200 Da.

#### 2.3 Liquid chromatography-mass spectrometry (Targeted Lipidomics)

Targeted lipidomics was performed on liver and plasma samples from 6 mice (from 3 virgin and 3 pregnant groups) and was additionally performed on samples from the separate WT and LXR DKO cohort (liver and serum samples from the groups at 14.5 dpc; liver, serum, placenta and fetal liver samples from the groups at 18.5 dpc).

Chromatographic separation of lipid and triglycerides was achieved using a Waters Acquity UPLC CSH C_18_ (50 mm × 2·1 mm, 1·7 μm) LC-column with a Shimadzu UPLC system (Shimadzu UK Limited, Milton Keynes, UK), at a flow rate of 0·5 mL/min maintained at 55 °C. Mass spectrometry detection was performed on a Thermo Exactive orbitrap mass spectrometer (Thermo Scientific, Hemel Hempstead, UK), which was operated in full scan mode from m/z 100–1800 Da (for singly charged species). To identify lipids, signal peaks were detected for the corresponding accurate mass at the correct retention time and were normalised to the total lipid/glyceride signal for that sample.

Signals data were collected across three runs: the first operating in positive ion and negative ionisation continuous switching mode, the second in a continuous negative ionisation mode switching between CID on and off, and the third was in a continuous positive ionisation mode switching between CID on and off. Runs 2 and 3 were used to determine the fatty acid composition of individual peaks in the chromatogram, thereby identifying the configuration of the lipid isoform(s) present.

#### 2.4 ^31^P NMR

Liver homogenates were combined into virgin (groups 1, 3 and 6) and pregnant (groups 2,5 and 8) pooled groups to give 5–10 mg of phospholipid per NMR sample. ^31^P NMR was run on the pooled samples using our recently described pipeline (82).

#### 2.5 Data Processing

Raw high-resolution mass-spectrometry data were processed using XCMS (www.bioconductor.org) and Peakpicker v2.0 (an in-house R script (83)). Theoretical lists of known species (by *m/z*) were used for both positive and negative ionisation modes (Supplementary Table S14). The correlation of signal intensity to concentration of the variable in liver and plasma QC samples (0.25, 0.50, 1.0) was used to identify which lipid signals were linearly proportional to concentration in the sample type and volume used (threshold for acceptance was a correlation of >0.75). Variables that deviated by more than 9 ppm, had a signal/noise ratio of <3 and had signals for fewer than 50% of samples were discarded, and zero values were interpreted as not measured. Relative abundance was then calculated by dividing each signal by the sum of signals for that sample and expressed per mille (‰). Final DI-MS datasets included 100 and 212 variables respectively in the positive and negative ionisation modes for plasma, and 141 and 315 variables respectively in the positive and negative ionisation modes for liver. All PC data was validated using signals from both ionisation modes and was presented using the positive ionisation mode data.

#### 2.6 Lipidomics Data Analysis

Positive and negative DI-MS datasets were analysed independently. Univariate analyses were conducted using Excel (Office 365) and multivariate analyses (MVA) were conducted using MetaboAnalyst 4.0 (84). Principal component analyses (PCA) were first performed to identify and exclude sample outliers based on 95% confidence intervals. Grouped lipid signals were compared between experimental groups using Two-way ANOVA with Sidak’s multiple comparisons. Individual lipid signals were compared between experimental groups using a combination of sparse Partial Least Squares-Discriminant Analyses (sPLS-DA, an unsupervised MVA) and Student’s *t*-tests, the p-values adjusted for multiple testing using a Bonferroni correction (0·05/sqrt(n)). Individual variables that passed both statistical tests, in at least two genotype-matched replicate comparisons, were regarded as the most important lipids describing the difference between conditions and were classified as candidate biomarkers (CBMs).

Phospholipid signals of interest were then selected for targeted analysis in the LC-MS/MS dataset to verify their signals and identified their associated fatty acids. *sn*-1/*sn*-2 fatty acid compositions were annotated using the most abundant isoform identified from the LC-MS/MS dataset. Triglyceride DI-MS signals whose FA residues have 54 or more carbons and 5 or more double bonds were expected to contain a LC-PUFA. Diglyceride DI-MS signals, which represent fragmented triglycerides, were expected to contain a LC-PUFA if they consisted of 38 or more carbons and 5 or more double bonds. For the independent WT and LXR DKO cohort, LC-MS/MS concentrations were compared between experimental groups using Bonferroni-adjusted Student’s *t*-tests.

### 3. PUFA Metabolites Analysis

#### 3.1 Lipid extraction

Liver samples were crushed with a FastPrep-24 Instrument (MP Biomedical, Fisher scientific SAS, Illkirch, France) in HBSS (Invitrogen, 200 μL) and deuterated internal standard mix (5 μL, 400 ng mL^−1^). After two crush cycles (6.5 ms^−1^, 30 s), an aliquot of the suspension (10 μL) was withdrawn for protein quantification and 0.3 mL of cold methanol added to the remaining material, which was then centrifuged (1016 × *g*, 15 min, 4 °C). The resulting supernatant was exposed to solid phase extraction using HLB plate (OASIS® HLB mg, 96- well plate, Waters, Saint-Quentin-en-Yvelines, France). Briefly, plates were conditioned with methanol (500 µL) and methanol-water (90:10, *v/v*, 500 µL). Samples were loaded at a flow rate of about 0.5 drop/s and, after complete loading, columns were washed with methanol-water (90:10, *v/v*, 500 µL). The columns were then dried under aspiration and lipids were recovered (methanol, 750 µL). The mixture was dried under a stream of N_2 (g)_ and samples were resuspended in methanol (140 μL) and transferred into a running vial (Macherey-Nagel, Hoerdt, France). The mixture was then dried and resuspended in methanol (10 µL) before injection onto the LC column.

#### 3.2 Liquid Chromatography/Tandem Mass Spectrometry data collection

6-keto-prostaglandin F1 alpha (6kPGF_1α_), thromboxane B2 (TxB_2_), Prostaglandin E2 (PGE_2_), 8-iso Prostaglandin A2 (8-isoPGA_2_), Prostaglandin E3 (PGE_3_), 15-Deoxy-Δ12,14- prostaglandin J2 (15d-PGJ_2_), Prostaglandin D2 (PGD_2_), Lipoxin A4 (LxA4), Lipoxin B4 (LxB4), Resolvin D1 (RvD1), Resolvin D2 (RvD2), Resolvin D5 (RvD5), 7-Maresin 1 (7- Mar1), Leukotriene B4 (LtB_4_), Leukotriene B5 (LtB_5_), Protectin Dx (PDx), 18- hydroxyeicosapentaenoic (18-HEPE), 5,6-dihydroxyeicosatetraenoic acid (5,6-DiHETE), 9- hydroxyoctadecadienoic acid (9-HODE), 13-hydroxyoctadecadienoic acid (13-HODE), 15- hydroxyeicosatetraenoic acid (15-HETE), 12-hydroxyeicosatetraenoic acid (12-HETE), 8- hydroxyeicosatetraenoic acid (8-HETE), 5-hydroxyeicosatetraenoic acid (5-HETE), 17- hydroxydocosahexaenoic acid (17-HDoHE), 14-hydroxydocosahexaenoic acid (14-HDoHE), 14,15-epoxyeicosatrienoic acid (14,15-EET), 11,12-epoxyeicosatrienoic acid (11,12-EET), 8,9-epoxyeicosatrienoic acid (8,9-EET), 5,6-epoxyeicosatrienoic acid (5,6-EET), 5- oxoeicosatetraenoic acid (5-oxoETE), Prostaglandin F2α, (PGF_2α_), 13- Hydroxyoctadecadienoic acid (13oxoODE), 9-hydroxyoctadecadienoic acid (9oxoODE), 9,10-dihydroxy-12-octadecenoic acid (9,10-DiHOME), 12,13-dihydroxy-12-octadecenoic acid (12,13-DiHOME), 9-hydroxy-10,12,15-octadecatrienoic acid (9-HOTrE), 13-hydroxy-9,11,15- octadecatrienoic acid (13-HOTrE), 9,10,13-trihydroxy-11-octadecenoic acid (10-TriHOME) and 9,12,13-trihydroxy-11E-octadecenoic acid (12-TriHOME), 3-hydroxydecanoic acid (C10- 3OH), 3-hydroxydodecanoic acid (C12-3OH), 3-hydroxytetradecanoic acid (C_14_-3OH), 3- hydroxyhexadecanoate (C_16_-3OH) and 3-hydroxyoctadecanoic acid (C_18_-3OH) were quantified. All the standards were purchased from Avanti® Polar Lipids (Millipore-SIGMA, St. Quentin Fallavier, France) and from Cayman Chemicals (Bertin Bioreagent, Montigny-le-Bretonneux, France). To simultaneously separate 45 lipids of interest and three deuterated internal standards (5-HETEd8, LxA4d4 and LtB4d4), LC-MS/MS analysis was performed on an ultrahigh-performance liquid chromatography system (UHPLC; Agilent LC1290 Infinity) coupled to an Agilent 6460 triple quadrupole MS (Agilent Technologies) equipped with electrospray ionization operating in negative mode. Reverse-phase UHPLC was performed using a Zorbax SB-C_18_ column (Agilent Technologies) with a gradient elution. The mobile phases consisted of water, acetonitrile (ACN), and formic acid (FA) [75:25:0.1 (*v/v/v*)] (solution A) and ACN and FA [100:0.1 (*v/v*)] (solution B). The linear gradient was as follows: 0% solution B at 0 min, 85% solution B at 8.5 min, 100% solution B at 9.5min, 100% solution B at 10.5 min, and 0% solution B at 12 min. The flow rate was 0.4 mL/min. The autosampler was set at 5 °C, and the injection volume was 5 µL. Data were acquired in multiple reaction monitoring (MRM) mode with optimized conditions (85).

#### 3.3 Data processing

Peak detection, integration, and quantitative analysis were performed with MassHunter Quantitative analysis software (Agilent Technologies). For each standard, calibration curves were built using 10 solutions at concentrations ranging from 0.95 to 500 ng/mL. A linear regression with a weight factor of 1/X was applied for each compound. The limit of detection (LOD) and the limit of quantification (LOQ) were determined for the 45 compounds using signal-to-noise ratios. The LOD corresponded to the lowest concentration leading to an S/N value >3, and LOQ corresponded to the lowest concentration leading to an S/N value >10. All values less than the LOQ were not considered. The concentration of LxB4, RvD1, RvD2, LTB_5_, 5,6-DiHETE, 14,15-EET and 11,12-EET did not reach the LOD in our samples. Blank samples were evaluated, and their injection showed no interference (no peak detected), during the analysis.

#### 3.4 PUFA metabolites data analysis

PUFA metabolites that best distinguish experimental groups were identified following the same CBM Discovery statistical workflow as described above.

### 4. Transcriptional Analysis

#### 4.1 Microarray dataset analyses

We re-analysed a previously published Affymetrix microarray of virgin and pregnant (14.5 dpc) mouse livers (n=4 per group) generated by Quinn and colleagues in 2019 (24). CEL files were downloaded from GEO using accession number GSE121202 and processed for QC and differential expression analysis using the Transcriptome Analysis Console (TAC) software (version 4.0.2; ThermoFisher). Genes were considered significantly differentially expressed if they exhibited at least a 1.25-fold difference in expression with a false discovery rate (FDR, based on the Benjamini–Hochberg procedure) of less than 0.05. Heatmaps were generated with R software packages (version 4.1.1) and volcano plots were generated using the EnhancedVolcano/Bioconductor package (version 3.15). Overlap analysis between upregulated differentially expressed genes and transcription factor targets was performed using the ChEA database in EnrichR (32).

#### 4.2 Liver mRNA expression

mRNA expression of genes of interest derived from the microarray analysis was measured in liver samples from selected experimental groups (virgin vs pregnancy analysis: groups 1, 2, 3 and 5; DLK1+ vs DLK1- analysis: groups 8, 7, 5 and 4) and in liver samples from the separate WT and LXR DKO cohort. For each analysis, all samples were processed together to avoid batch effects. Frozen liver samples were homogenized in TRIzol^TM^ (Thermo Fisher: Waltham, MA, USA) and kept at −80°C until RNA extraction following the manufacturer’s protocol. RNA samples were treated with DNase I (M0303, New England Biolabs (NEB)) and sodium acetate precipitated. RNA integrity was assessed by the identification of intact 18S and 28S rRNA bands after agarose gel electrophoresis and purity was confirmed by 260/280 nm absorbance ratios calculated by a NanoDrop spectrophotometer. Complementary DNA (cDNA) was obtained by reverse transcription (RT) using 1 μg of RNA and Moloney murine leukaemia virus (M-MuLV) reverse transcriptase (M0253, NEB), using the first strand cDNA synthesis standard protocol with random primers (S1330, NEB), 2.5 mM dNTPs (N0446, NEB) and RNase inhibitors (M0314, NEB).

Quantitative real-time PCR (RT-qPCR) assays were performed using SYBR Green qRT-PCR master mix (QuantiNova SYBR Green PCR Kit, Qiagen) with primers designed using the Primer3 software (primer3.ut.ee) or using previously reported sequences (4,86–89) (see Supplementary Table S15 for complete list) and ordered from Millipore-SIGMA. cDNA samples were diluted 20x and quantification was performed using the relative standard curve method. For the virgin vs pregnancy analysis, target gene expression was normalized to the average expression of *Hprt*, *α-tubulin* and *Tbp* and compared between group-pairs by Mann–Whitney U tests. Multiple correlation coefficients with Bonferroni corrected p-values were calculated for virgin groups (groups 1 and 3) and pregnant groups (groups 2 and 5) to correlate the expression pattern of multiple genes within each condition. For the DLK1+ vs DLK1- analysis, genes were normalised to the average expression of *Hprt* and *α-tubulin.* Normalised gene expression was compared between all four groups in the DLK1 analysis by one-way ANOVA with Tukey’s multiple comparison post hoc tests. For the WT vs LXR DKO analysis, gene expression was normalized to the average expression of *Hprt*, *α-tubulin* and *Tbp* and compared by two-way ANOVA with Šídák’s multiple comparisons.

### 5. Immunohistochemistry

Fresh livers cut into a ∼5mm wide sagittal slice at the level of the gall bladder then were fixed with Neutral Buffered Formalin (Millipore-SIGMA, HT501128) overnight at 4°C and dehydrated through an increasing ethanol series the following day. Samples dehydrated to 100% ethanol were then incubated with Histoclear II (National Diagnostics, HS202) (2 x 2hr incubations), followed by 2 x 2-hour incubations at 65°C with Histosec® (1.15161.2504, VWR). 5μm histological sections were cut using a Thermo HM325 microtome, mounted on Menzel-Gläser Superfrost®Plus slides (Thermo Scientific, J1810AMNZ) and used for immunofluorescence (IF).

For double IF, histological sections were rehydrated then antigen unmasking was achieved by incubation in boiling Tris-EDTA buffer pH 9 [10 mM Tris Base, 1 mM EDTA] for 20 mins. Histological sections were incubated overnight at 4°C with the primary antibodies (rabbit anti-human COX1, ab133319, abcam, 1:200; rat anti-mouse Endomucin, sc-5495 Santa Cruz, 1:200), then incubated with Texas Red-conjugated goat anti-Rat (TI-9400 Vector Labs, 1:300) and goat α-rabbit DyLight® 488 (DI1488, Vector Laboratories, 1:300) for 1hr at RT, and mounted using VECTASHIELD® Antifade Mounting Medium with DAPI (H-1200-10, Vector Laboratories). Isotype controls for each antibody showed no staining under identical conditions.

### 6. Western Blot

Frozen liver samples were each lysed in 200 µl Pierce^TM^ RIPA lysis buffer (Thermo Fisher #89900) with 1% Halt^TM^ protease and phosphatase inhibitor (Thermo Scientific #78442). Protein concentrations were determined using the Pierce^TM^ BCA protein assay kit (Thermo Scientific #23227) and all lysates were diluted to 4 µg/µl in Laemmli buffer (National Diagnostics EC-886-10). Proteins were denatured at 95°C for 5 minutes and run on a Bolt™ 4-12% Bis-Tris polyacrylamide gel (Thermo Scientific #NW04125BOX). Proteins were transferred to a nitrocellulose membrane (Invitrogen #PB3310) using the Power Blotter– Semi-dry Transfer System (Invitrogen #PB0013). Membranes were blocked with 5% dairy milk in TBST (100 mM Tris (pH7.5), 400mM NaCl, 0.05% Tween-20) and incubated in anti-PEMT rabbit polyclonal antibody (1:1000, Bio-techne #NBP1-59580) or anti-α-TUBULIN mouse monoclonal antibody (1:10,000, Millipore-SIGMA #T5168) in 5% dairy milk, overnight at room temperature. Membranes were then washed in TBST and incubated with the secondary HRP-conjugated antibodies: goat anti-rabbit (1∶1000, P0448, DAKO) or polyclonal anti-mouse (1∶2000, P0447, DAKO) in 5% dairy milk for 1 hour at room temperature. Following washing in TBST, membranes were treated with Pierce^TM^ ECL substrate (Thermo Scientific #32132) and chemiluminescence was detected using the iBright FL1000 Imaging System (Invitrogen).

### 7. Statistical Analyses

For all comparisons between experimental cohorts, significance was only considered if identified in at least two genotype-matched replicate comparisons. Unless otherwise stated, all statistical tests were performed using the GraphPad Prism Software, version 8.4.3 for Windows (www.graphpad.com).

## Notes

### Competing Interest Statement

The authors have declared no competing interest.

