## Supplementary Figures for "A co-ordinated transcriptional programme in the maternal liver supplies LC-PUFAs to the conceptus using phospholipids"

**Supplementary Figure 1**

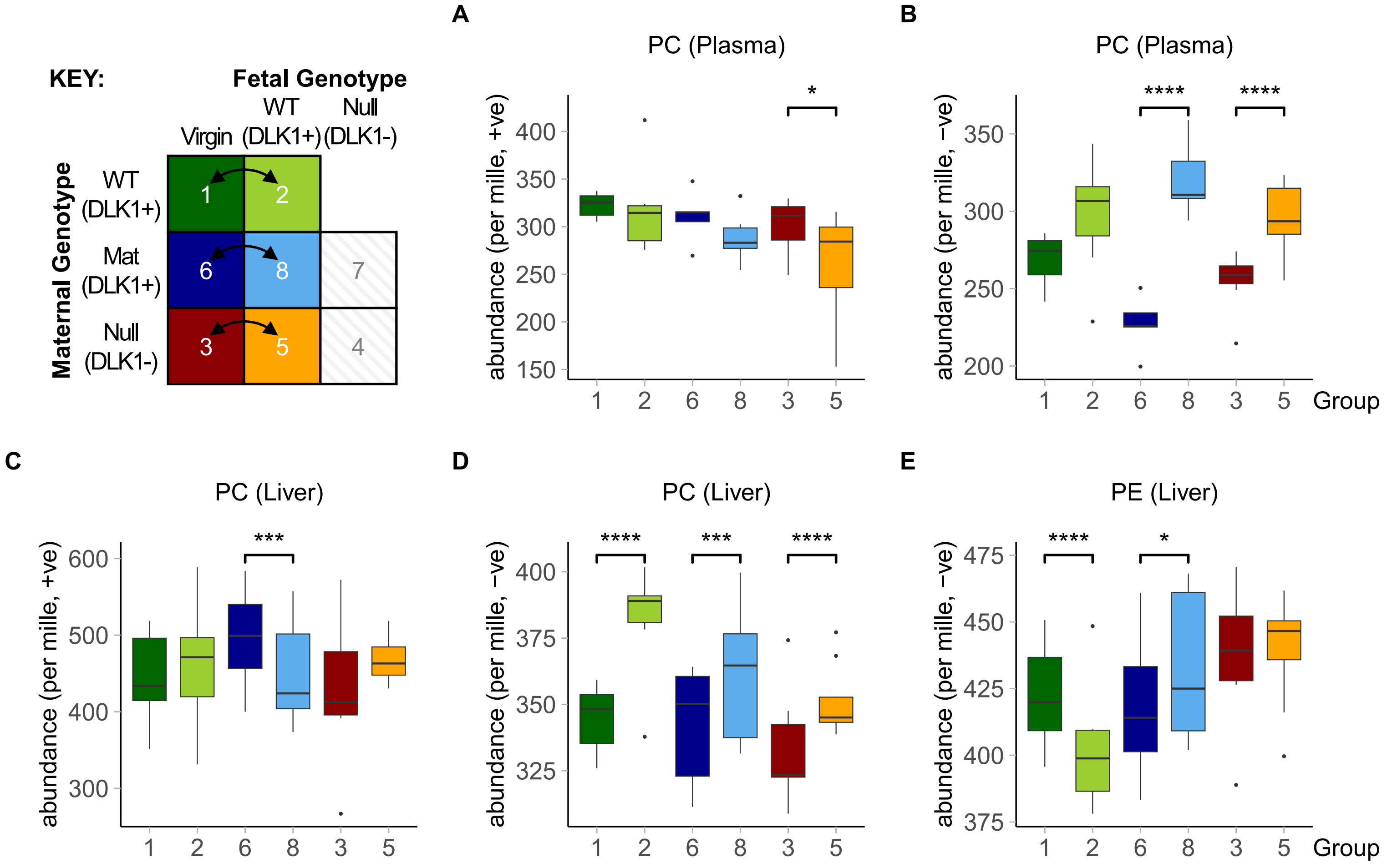

**Supplementary Figure 2**

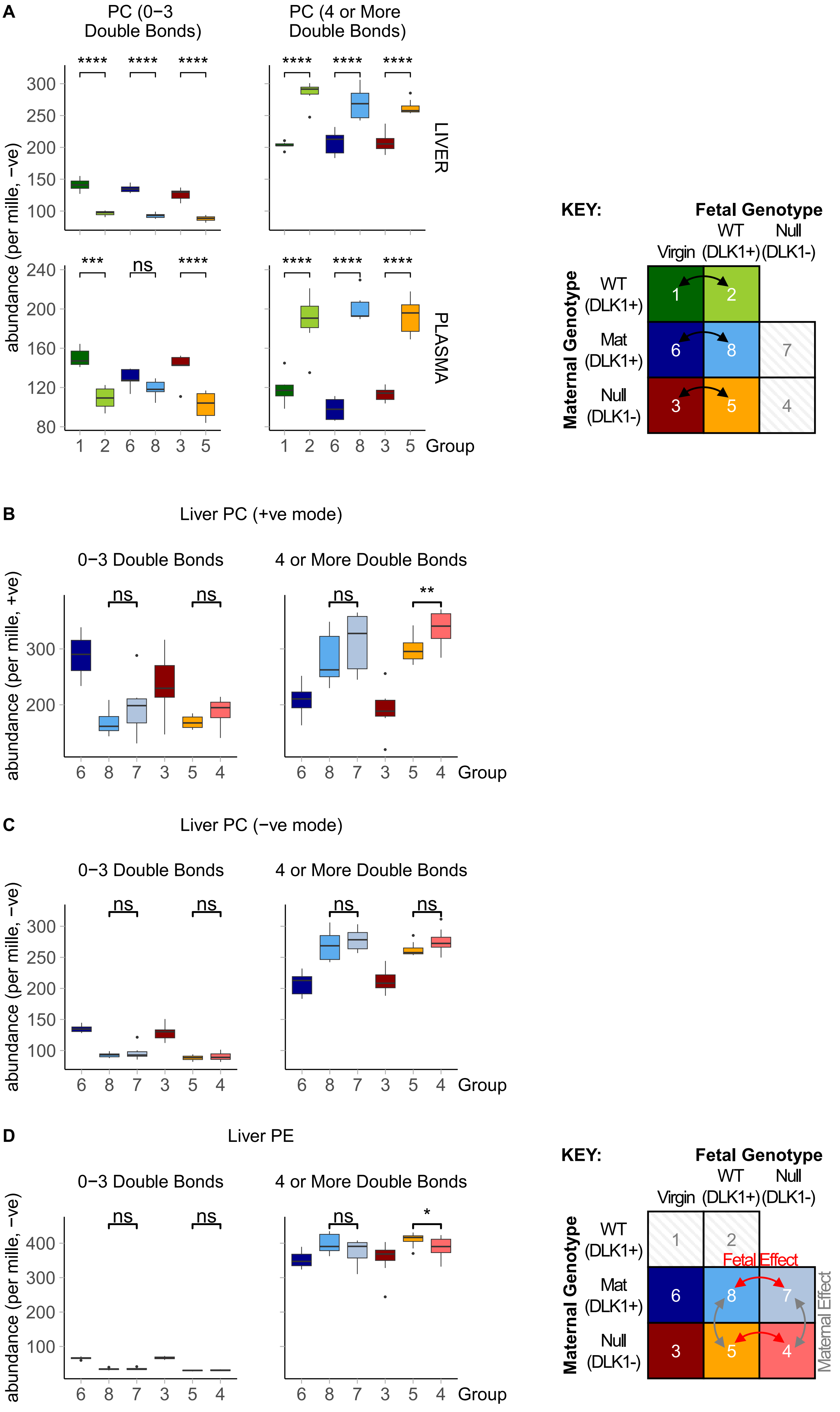

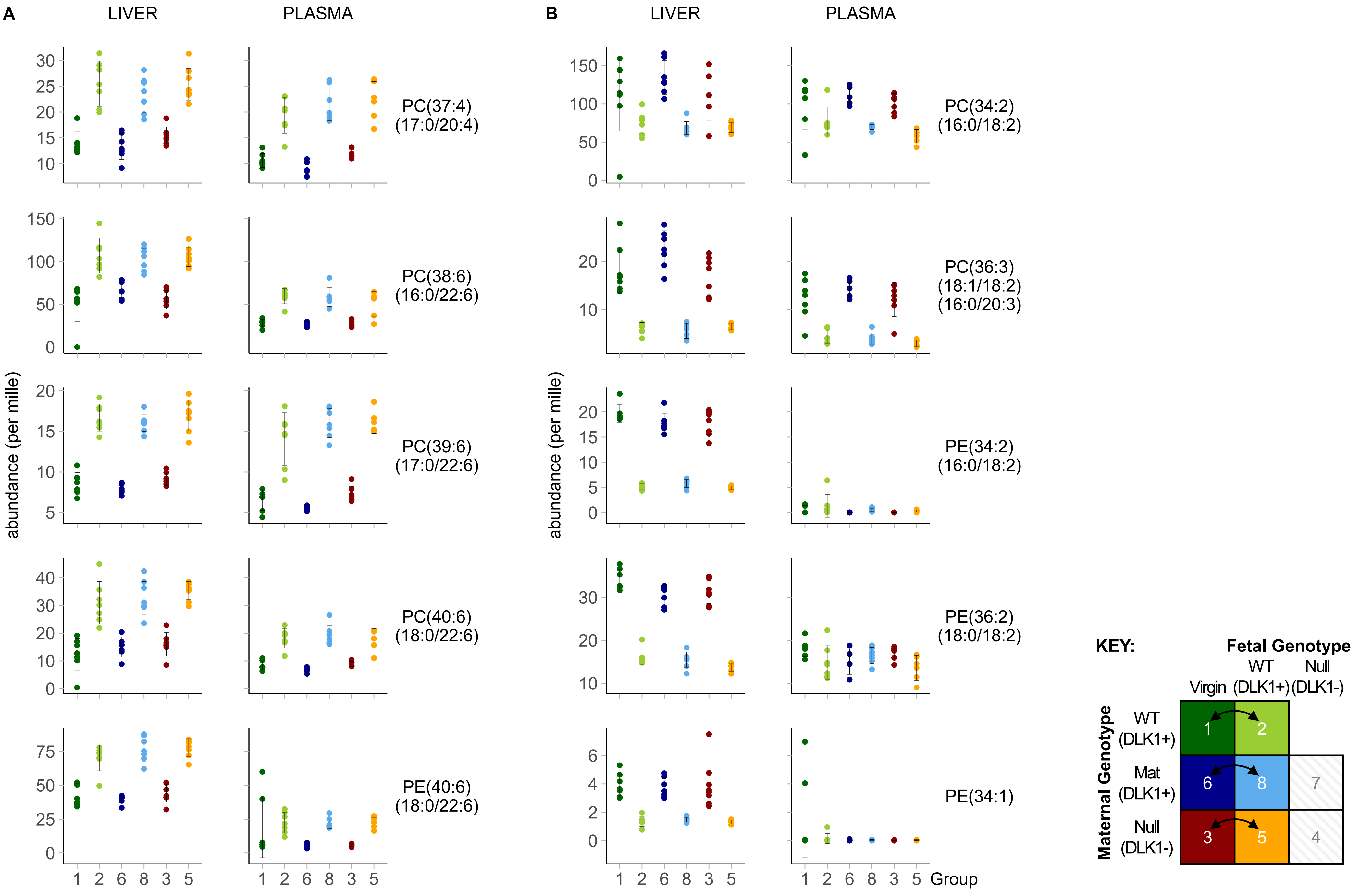
A
LIVER
PLASMA
B
LIVER
PLASMA
abundance (per mille)
PC(37:4)
(17:0/20:4)
PC(38:6)
(16:0/22:6)
PC(39:6)
(17:0/22:6)
PC(40:6)
(18:0/22:6)
PE(40:6)
(18:0/22:6)
PC(34:2)
(16:0/18:2)
PC(36:3)
(18:1/18:2)
(16:0/20:3)
PE(34:2)
(16:0/18:2)
PE(36:2)
(18:0/18:2)
PE(34:1)
Group
KEY:
Fetal Genotype
Virgin
WT
(DLK1+)
Null
(DLK1-)
Maternal Genotype
WT
(DLK1+)
Mat
(DLK1+)
Null
(DLK1-)

**Supplementary Figure 4**

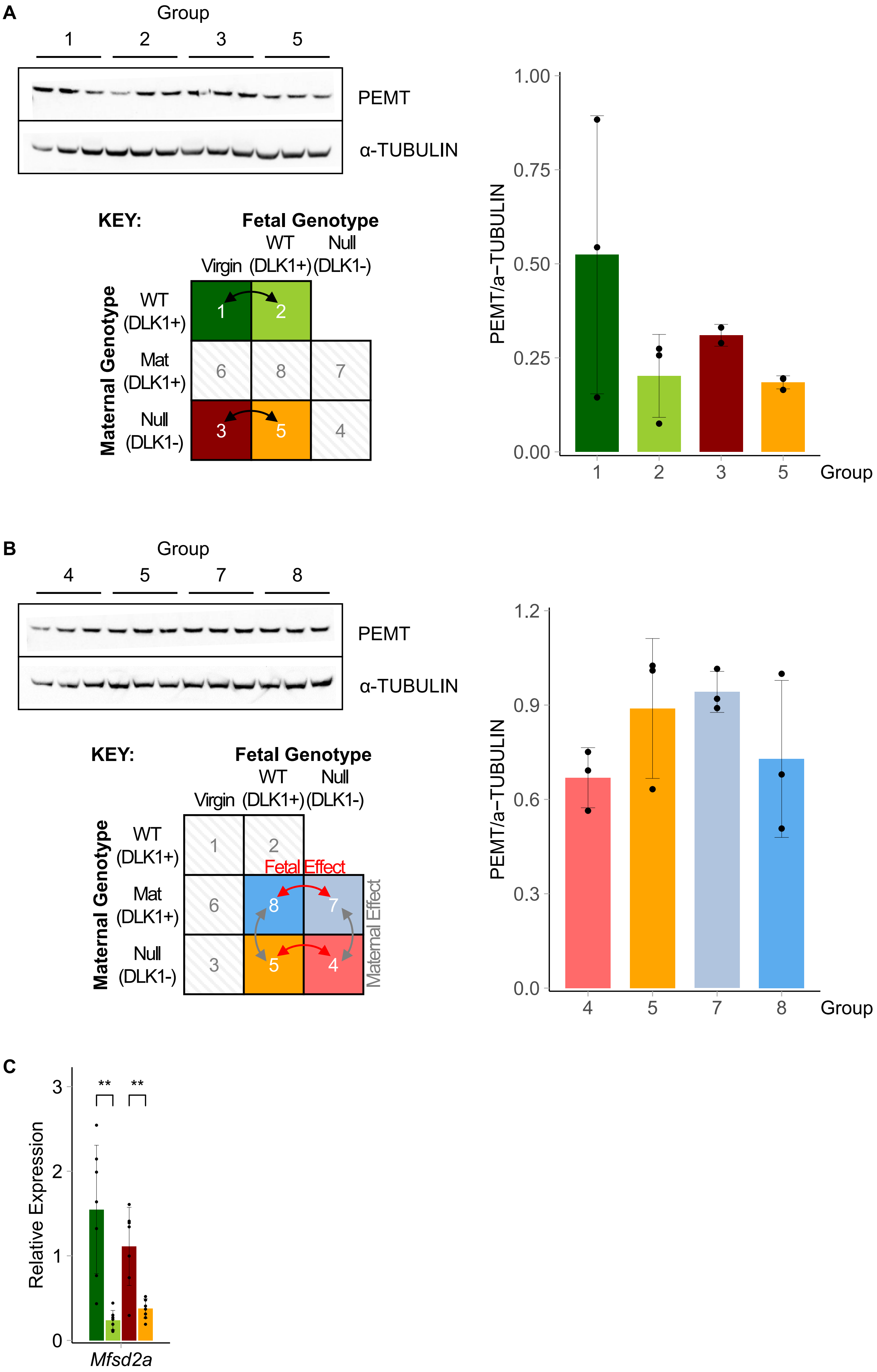

**Supplementary Figure 5**

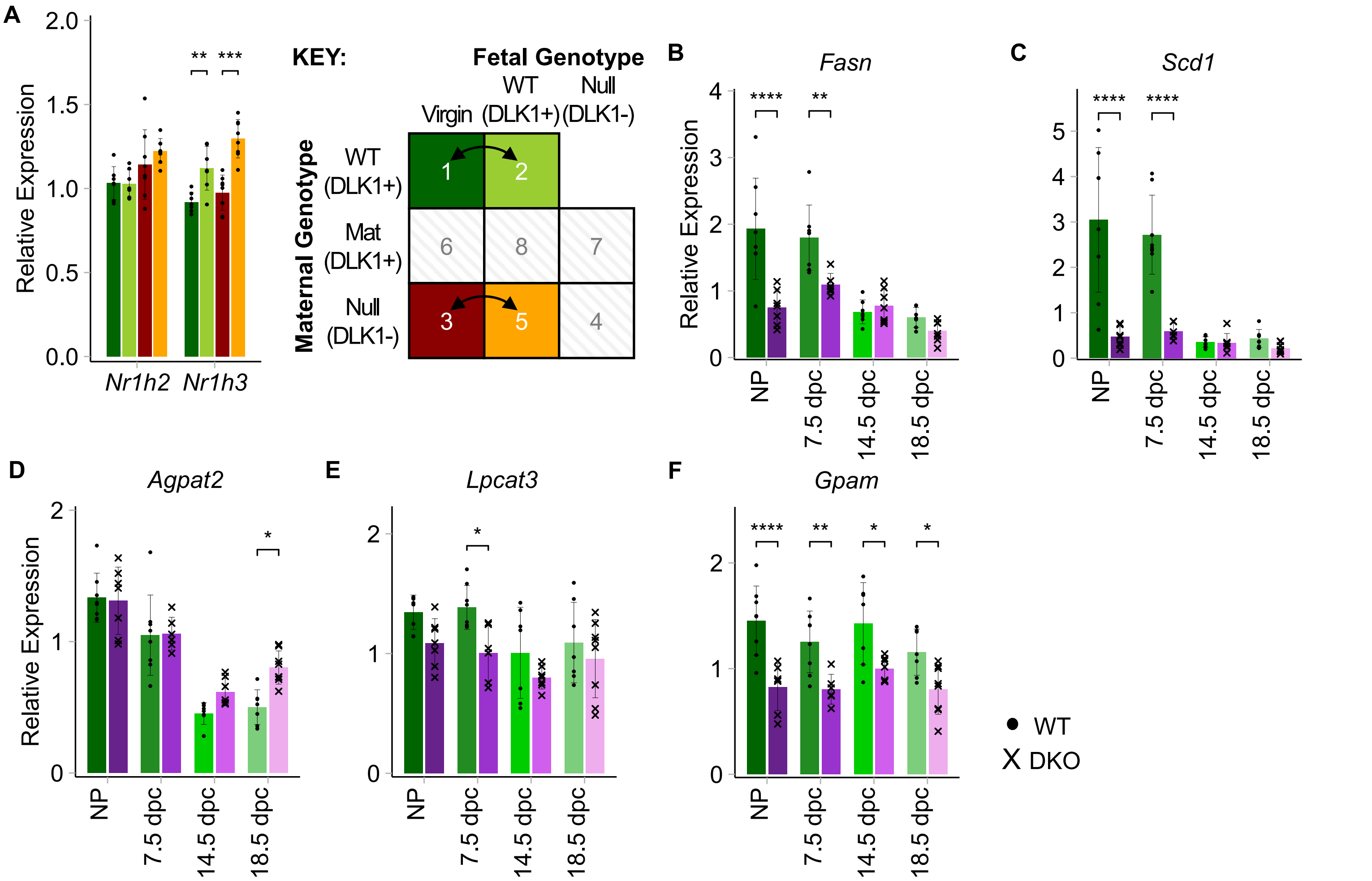

**Figure S1:** Grouped relative abundance of PC in plasma **(A-B)** and liver **(C-D)** and hepatic PE **(E)** in three genotype-matched virgin vs pregnant (15.5 dpc) replicate group comparisons. PC data is shown from both positive and negative ionisation modes. Data is presented as boxplots with whiskers showing 1.5*IQR and outliers plotted individually. Two-way ANOVA with Sidak's multiple comparisons tests were performed to determine significant class shifts between experimental groups (* p-value <0.05; ** p-value <0.01; *** p-value <0.001; **** p-value <0.0001). Statistical tests were performed independently per ionisation mode and per replicate comparison. n=5-8 per group. PC, phosphatidylcholine; PE, phosphatidylethanolamine.

**Figure S2: (A)** Grouped relative abundance of PC lipids measured in the negative ionisation mode that contain fatty acids with a combined total of three or fewer double bonds (left) or four or more double bonds (right) in the liver and plasma of virgin and pregnant (15.5 dpc) groups. **(B-D)** Grouped relative abundance of PC **(B-C)** and PE **(D)** lipids that contain fatty acids with a combined total of three or fewer double bonds (left) or four or more double bonds (right) in livers of pregnant mice that lack fetal or maternal-derived DLK1 protein. Significance was only considered if identified in at least two genotype-matched replicate group comparisons. Grouped abundance data is presented as boxplots (whiskers showing 1.5*IQR and outliers plotted individually) and two-way ANOVA with Sidak's multiple comparisons were performed for each genotype-matched comparison (* p-value <0.05; ** p-value <0.01; *** p-value <0.001; **** p-value <0.0001). All statistical tests were performed independently per ionisation mode and per replicate comparison. n=5-8 per group. PC, phosphatidylcholine; PE, phosphatidylethanolamine.

**Figure S3:** PC or PE lipids of interest that contain an *sn*-2 fatty acid with four or more double bonds **(A)** and three or fewer double bonds **(B)** that were identified as candidate biomarkers (CBMs) associated with pregnancy. CBMs are classified as lipids that passed both Bonferroni-adjusted *t*-tests (liver threshold, p = 0.00234; plasma threshold, p = 0.00283) and sparse partial least squares discriminant analysis in at least two genotype-matched replicate virgin vs pregnant (15.5 dpc) comparisons (see Supplementary Table S4 for full CBM list). CBM data is shown as relative abundance per ionisation mode with ± SD error bars. *sn*-1/*sn*-2 fatty acid compositions were assigned using the most abundant isoform identified from targeted LC-MS/MS analysis in plasma and liver (Supplementary Table S6). n=5-8 per group. PC, phosphatidylcholine; PE, phosphatidylethanolamine.

**Figure S4: (A-B)** Western blotting of PEMT protein levels from liver lysates of two genotype-matched virgin vs pregnant (15.5 dpc) group comparisons **(A)** and in the four pregnant groups used to assess the effect of maternal-derived DLK1 protein or fetal-derived DLK1 protein in pregnancy **(B)**. PEMT bands are normalised to the housekeeping protein α-tubulin. n = 3 biological replicates/group. **(C)** Real-time quantitative PCR (RT-qPCR) of lyso-phospholipid transporter, *Mfsd2a,* in livers from two genotype-matched virgin vs pregnant group comparisons (n=7-8/group). Data was normalised to housekeeping gene expression (*Tuba1*, *Tbp* and *Hprt*) and is shown as mean relative expression ± SD. Groups were called significantly different by Mann-Whitney U test (* p-value <0.05; ** p-value <0.01; *** p-value <0.001).

**Figure S5: (A)** Real-time quantitative PCR (RT-qPCR) analysis of liver X receptor (LXR) isoforms, LXRα (*Nr1h3*) and LXRβ (*Nr1h2*), in livers from two genotype-matched virgin vs pregnant (15.5 dpc) group comparisons (n=7-8/group). **(B-F)** RT-qPCR analysis of classical lipogenic LXR target genes **(B-C)** and additional LC-PUFA-phospholipid biosynthetic genes **(D-F)** in an independent cohort of wild-type (WT) and *Lxrab*$^{-/-}$ (LXR double knockout (DKO)) mice at non-pregnant and various gestational timepoints (n=6-8/group). RT-qPCR data was normalised to housekeeping gene expression (*Tuba1*, *Tbp* and *Hprt*) and is shown as mean relative expression ± SD. Virgin vs pregnant groups were compared by Mann-Whitney U test, WT vs LXR DKO groups were compared by two-way ANOVA with Šídák's multiple comparison (* p-value <0.05; ** p-value <0.01; *** p-value <0.001; **** p-value <0.0001).
